# Characterization of non-canonical G beta-like protein FvGbb2 and its relationship with heterotrimeric G proteins in *Fusarium verticillioides*

**DOI:** 10.1101/781484

**Authors:** Huijuan Yan, Won Bo Shim

**Affiliations:** Department of Plant Pathology and Microbiology Texas A&M University, College Station 77843, TX, USA

## Abstract

*Fusarium verticillioides* is a fungal pathogen that is responsible for maize ear rot and stalk rot diseases worldwide. The fungus also produces carcinogenic mycotoxins, fumonisins, on infested maize. Unfortunately, we still lack clear understanding of how the pathogen responds to host and environmental stimuli to trigger fumonisin biosynthesis. The heterotrimeric G protein complex, consisting of canonical Gα, Gβ, and G*γ* subunits, is involved in transducing signals from external stimuli to regulate downstream signal transduction pathways. Previously, we demonstrated that Gβ protein FvGbb1 has direct impact on fumonisin regulation but no other physiological aspects in *F. verticillioides*. In this study, we identified and characterized a putative receptor for activated C kinase 1 (RACK1) homolog FvGbb2 as a putative Gβ-like protein in *F. verticillioides.* The mutant exhibited severe defects not only in fumonisin biosynthesis but also vegetative growth and conidiation. FvGbb2 was positively associated with carbon source utilization and stress agents but negatively regulated general amino acid control. While FvGbb2 does not interact with canonical G protein subunits, it may interact with diverse proteins in the cytoplasm to regulate vegetative growth, virulence, fumonisin biosynthesis, and stress response in *F. verticillioides*.

## Introduction

*Fusarium verticillioides* (teleomorph: *Gibberella moniliformis* Wineland) is a widely distributed fungal pathogen associated with every stage of the maize life cycle, from seed germination to harvest and post-harvest storage (Blacutt et al., 2018). The most common maize diseases caused by this fungus are stalk rot, ear rot and seedling blight. Importantly, *F. verticillioides* produces a variety of mycotoxins including fusaric acid, fusarins, and fumonisins. Fumonisin B1 is the most abundant and toxic form among fumonisins, a group of polyketide-derived secondary metabolites, that contaminate cereals and grains, leading to health and food safety risks for humans and animals. Note that a wide variety of secondary metabolites are synthesized in microbes, plants, and even certain marine animals. Polyketide synthases (PKSs) and non-ribosomal peptide synthases (NRPSs) are mainly responsible for the biosynthesis of diverse secondary metabolites. To date, 16 PKSs and 16 NRPSs are identified in *F. verticillioides* genome (Hansen et al., 2015). Particularly *PKS11* (also known as *FUM1*) is the essential PKS for initiating fumonisin biosynthesis. While some PKS genes have been characterized in *F. verticillioides* secondary metabolism, *e.g*. *PKS3* (*PGL1*) is required for the dark violet pigment production in perithecia and *PKS10* contributes to fusarin biosynthesis, the majority of PKS gene functions remain unclear (Hansen et al., 2015).

Furthermore, these secondary metabolites are not required for normal fungal growth, survival, or virulence but are known to provide physiological benefits, including protection against UV damage, defense against abiotic and biotic stresses (Ma et al., 2013; Keller, 2018). It is also important to note that there is still a significant gap in our knowledge on how these secondary metabolites, including fumonisins, are regulated for biosynthesis.

The canonical heterotrimeric G protein complex, which consists of *α*, *β*, and *γ* subunits (Neves et al., 2002), is critical for sensing environmental stimuli, including the pheromones, nutrients and host infection. When activated by their cognate ligands, seven transmembrane domain receptors, *i.e.* G protein-coupled receptors (GPCRs), will go through a conformational change. This leads to the exchange of GTP for GDP in the Gα subunit triggering the dissociation of G protein complex (Xue et al., 2008). Both the Gα subunit and the Gβγ dimer are responsible for initiating downstream signaling pathway including cAMP and MAPK kinase pathways. Multiple Gα subunits are identified in filamentous fungi, while only one canonical Gβ subunit is found in fungal species (Li et al., 2007). Gβ subunit is best studied WD40-repeat protein. In addition to harboring seven WD40-repeat domains, Gβ subunit also contains a short coiled-coil motif in the N-terminus region responsible for interacting with G*γ* subunit (Zeller et al., 2007). In addition to the Gβ subunit, 12 out of 55 WD proteins have seven WD40-repeat domains in *S. cerevisiae*, but only one Gβ and one Gβ-like protein were identified (Smith et al., 1999). Studies suggest that Gβ-like protein is associated with diverse physiological functions in fungi (Zhang et al., 2016).

Putative Gβ-like protein RACK1 was first identified from a chicken liver cDNA library. Later, RACK1 was also identified as a receptor protein for Activated Protein Kinase C (PKC) (Mochlyrosen et al., 1991; Ron et al., 1994). In *Aspergillus fumigatus*, the deletion of RACK1 homolog CpcB led to defects in hyphal growth, virulence and gliotoxin biosynthesis (Kong et al., 2013; Cai et al., 2015). In rice blast pathogen *Magnaporthe oryzae*, RACK1 homolog Mip11 was identified as a Mst50- and MoRgs7-binding protein, which is critical in cAMP signaling and plant infection (Li et al., 2017; Yin et al., 2018). RACK1 homologs were shown to interact directly with one of Gα subunits in multiple fungi, including *S. cerevisiae*, *Cryptococcus neoformans* and *M. oryzae* (Palmer et al., 2006; Zeller et al., 2007). Furthermore, a recent study in *Arabidopsis thaliana* demonstrated that RACK1 actually serves as a scaffolding protein for Gβ subunit and a downstream MAPK cascade component involved in plant immunity (Cheng et al., 2015). If RACK1 were to be recognized as a Gβ-like protein in *F. verticillioides*, one question we can ask is whether this protein physically interacts with heterotrimeric G protein subunits.

In *F. graminearum*, the deletion mutant of Gβ protein GzGpb1 facilitated ZEA and DON production but was defective in the hyphal growth and pathogenicity (Yu et al., 2008). Our previous study demonstrated that Gβ subunit FvGbb1 positively regulated FB_1_ biosynthesis, but otherwise showed very limited roles in *F. verticillioides* vegetative growth and virulence (Sagaram and Shim, 2007). Significantly, as described earlier, Gβ-like subunit RACK1 is involved in diverse functions in multiple eukaryotic organisms. The outcomes from our FvGbb1 study led us to hypothesize that an alternative non-conventional Gβ-like subunit may compensate for the main Gβ subunit functions in *F. verticillioides*. In this study, we characterized the functional role of putative Gβ-like protein FvGbb2, a RACK1 homolog, in *F. verticillioides* development and secondary metabolisms. Additionally, we investigated the relationship between FvGbb2 and canonical heterotrimeric G protein components.

## Results

### *Identification of the RACK1 homolog in* F. verticillioides

We used Asc1 protein sequence from the *S. cerevisiae* genome database (http://www.yeastgenome.org/) to search for RACK1 homolog in *F. verticillioides* database using BLASTP algorithm. The search led to the identification of FVEG_02582, which was designated as FvGbb2. The *FvGBB2* gene is 1,465 bp in length with 3 predicted introns, encoding a 316- amino-acid protein. To identify the FvGbb2 orthologs in other species, *F. verticillioides* FvGbb2 amino acid sequence was used for a BLAST query. Sequence alignment indicates that FvGbb2 protein shared a high amino acid identity with its orthologs (Fig. S1). Both FvGbb2 and FvGbb1 are predicted to contain seven WD-40 repeat motifs, while FvGbb1 has an extra coiled-coil domain. FvGbb2 did not share high protein similarity with FvGbb1, but this is not surprising when you consider structural diversity of WD-40 repeat motifs and the difficulty associated with predicting secondary structure information (Wang et al., 2016). But, to further study FvGbb2 against canonical Gβ subunits in fungi, amino acid sequences of candidate proteins were aligned with ClustalW software (Larkin et al., 2007), and the phylogenetic tree was built with bootstrap 1000 replicates using MEGA 7 (Kumar et al., 2016). Domain information was retrieved from SMART and presented by EvolView (Letunic et al., 2015; He et al., 2016; Letunic and Bork, 2018) (Fig. 1). Phylogenetic tree analysis suggested that FvGbb2 and its homologs are distinct from the canonical Gβ subunit in various species.

**Fig. 1.**
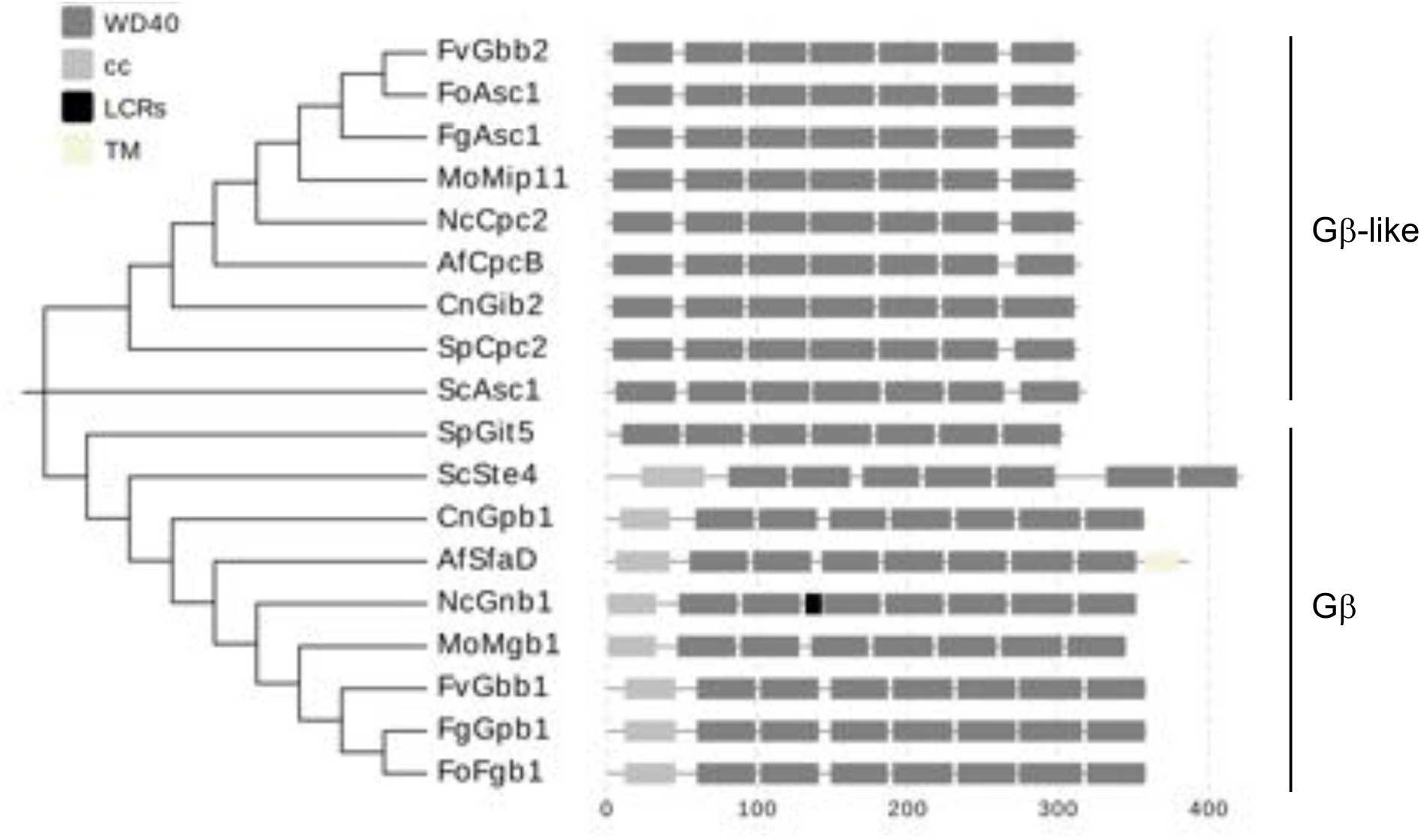
Phylogenetic and domain analysis of Gβ-like and Gβ proteins in multiple fungi. Protein names with fungal species, NCBI locus tag, and protein similarity against FvGbb2 used in the figure are; FvGbb2 (*F. verticillioides*, FVEG_02582, 100% identity), FoAsc1 (*F. oxysporum* f. sp. *lycopersici* 4287, FOXG_05557, 100% identity), FgAsc1 (*F. graminearum,* FGSG_09870, 99% identity), MoMip11 (*Magnaporthe oryzae,* MGG_04719, 97% identity), NcCpc2 (*Neurospora crassa* OR74A, NCU05810, 95% identity), AfCpcB (*Aspergillus fumigatus* A1163, AFUB_070060, 90% identity), CnGib2 (*Cryptococcus neoformans var. grubii H99*, CNAG_05465, 73% identity), SpCpc2 (*Schizosaccharomyces pombe,* SPAC6B12.15, 70% identity), ScAsc1 (*Saccharomyces cerevisiae*, YMR116C, 59% identity), SpGit5 (*S. pombe*, SPBC32H8.07, 22% identity), ScSte4 (*S. cerevisiae*, YOR212W, 26% identity), CnGpb1 (*C. neoformans var. grubii H99*, CNAG 01262, 22% identity), AfSfaD (*A. fumigatus* A1163, AFUB_059800, 23% identity), NcGnb1 (*N. crassa* OR74A, NCU00440, 23% identity), MoMgb1 (*M. oryzae*, MGG_05201, 22% identity), FvGbb1 (*F. verticillioides*, FVEG_10291, 22% identity), FgGpb1 (*F. graminearum*, FGSG_04104.1, 27% identity), FoFgb1 (*F. oxysporum* f. sp*. lycopersici* 4287, FOXG_11532, 22% identity).

### ΔFvgbb2 had severe defects in vegetative growth and conidia on agar plates but showed more vigorous mycelial growth in liquid YEPD medium

To study the functional role of FvGbb2, we generated a null mutant of *FvGBB2* (ΔFvgbb2) by replacing the gene with the hygromycin-resistance gene (Fig. S2A). *FvGBB2* transcript in the mutant was tested by qPCR, and the complete loss of *FvGBB2* expression indicated successful gene deletion (Fig. S2D). The ΔFvgbb2 mutant grew significantly slower and less fluffy compared to the wild type (WT) on V8, 0.2xPDA, myro and YEPD agar plates (Fig. 2A). In addition, 7-day-old YEPD liquid cultures of double deletion mutants ΔFvgbb2-gbb1, ΔFvgbb2-gpa2 showed extremely high levels of undetermined dark pigment (Fig. 2A). Additionally, when examined under the microscope, ΔFvgbb2 exhibited shorter but noticeable hyper-branching hyphae (Fig. 2B). Gene complementation strains ΔFvgbb2-Com and ΔFvgbb2-GFP showed full recovery of the growth defects in the ΔFvgbb2 strain. These results indicated that Fvgbb2 is important for vegetative growth in *F. verticillioides*. Moreover, we measured the number of conidia after 8 days incubation on V8 plates at room temperature. ΔFvgbb2 exhibited dramatic inhibition of conidia production and germination (Fig. 3A and 3C). Interestingly, we found that ΔFvgbb2 mycelium mass (fresh weight) in YEPD liquid media was significantly higher than WT (Fig. 3B). To further understand the molecular basis of these defects, we studied expression levels of conidia-related genes including *FvBRLA*, *FvWETA*, *FvABAA* and *FvSTUA.* Total RNA was extracted from WT and mutant mycelia collected from YEPD liquid cultures incubated for 20 h with agitation (150 rpm). Transcript levels of conidia-related genes were all significantly down-regulated in ΔFvgbb2 in myro and YEPD liquid media (Fig. 3D and S3A). We also tested whether FvGbb2 is essential for sexual reproduction, but all mutant strains were capable of perithecia production (Fig. S3B).

**Fig. 2.**
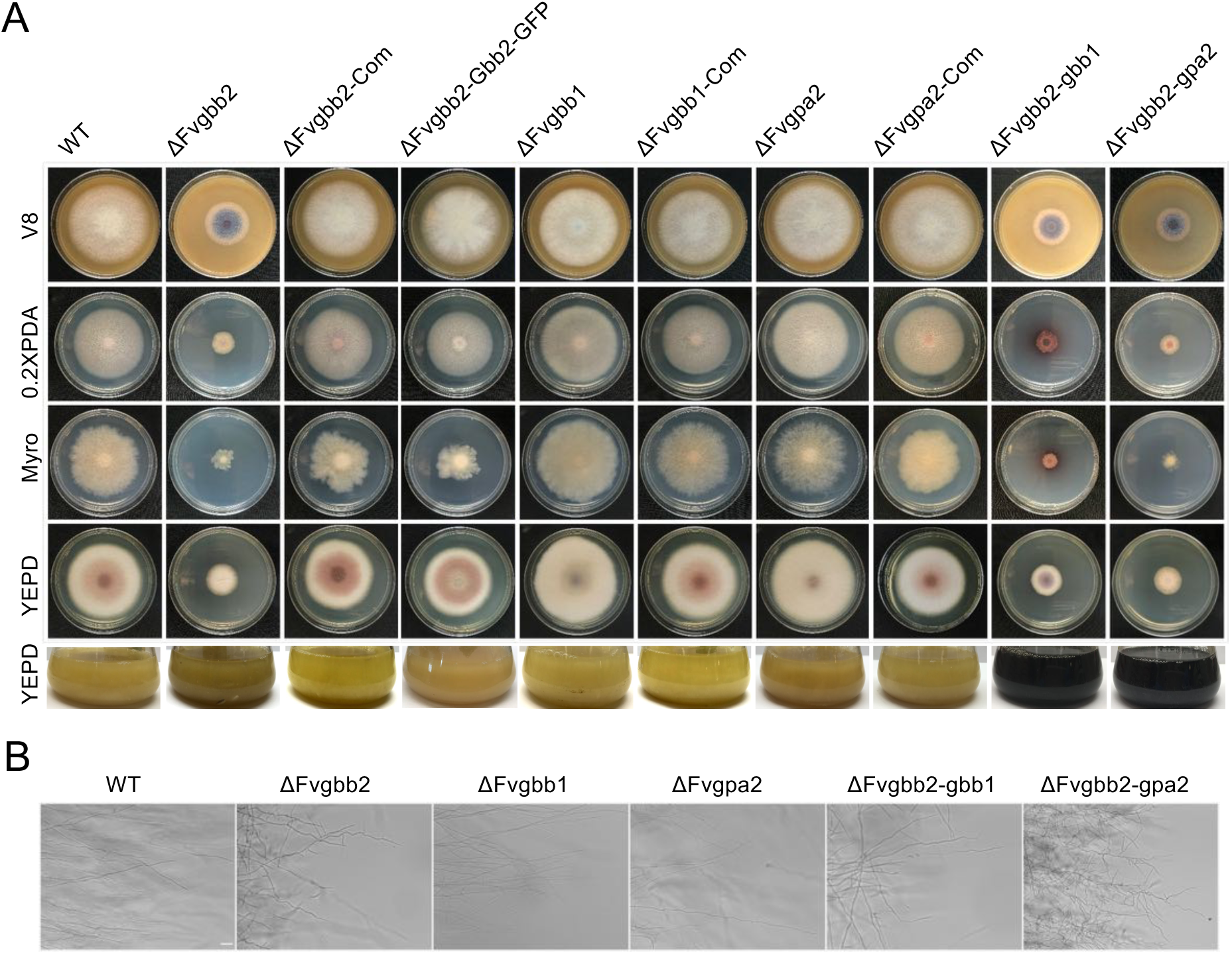
Mycelial growth and morphology of ΔFvgbb2, ΔFvgbb1, ΔFvgpa2, ΔFvgbb2-gbb1, and ΔFvgbb2-gpa2 mutants. (A) Colonies of the wild-type (WT), ΔFvgbb2, ΔFvgbb2-Com, ΔFvgbb2-Gbb2-GFP, ΔFvgbb1, ΔFvgbb1-Com, ΔFvgpa2, ΔFvgpa2-Com, ΔFvgbb2-gbb1, ΔFvgbb2-gpa2 mutants grown on V8 agar, 0.2xPDA, myro agar, YEPD agar plates for 8 days at room temperature. Strains were also cultured in YEPD liquid medium for 7 days with agitation at 150 rpm. (B) Hyphal growth and branching were examined on 0.2xPDA agar after 3 days of incubation at room temperature. Bar = 100 μm

**Fig. 3.**
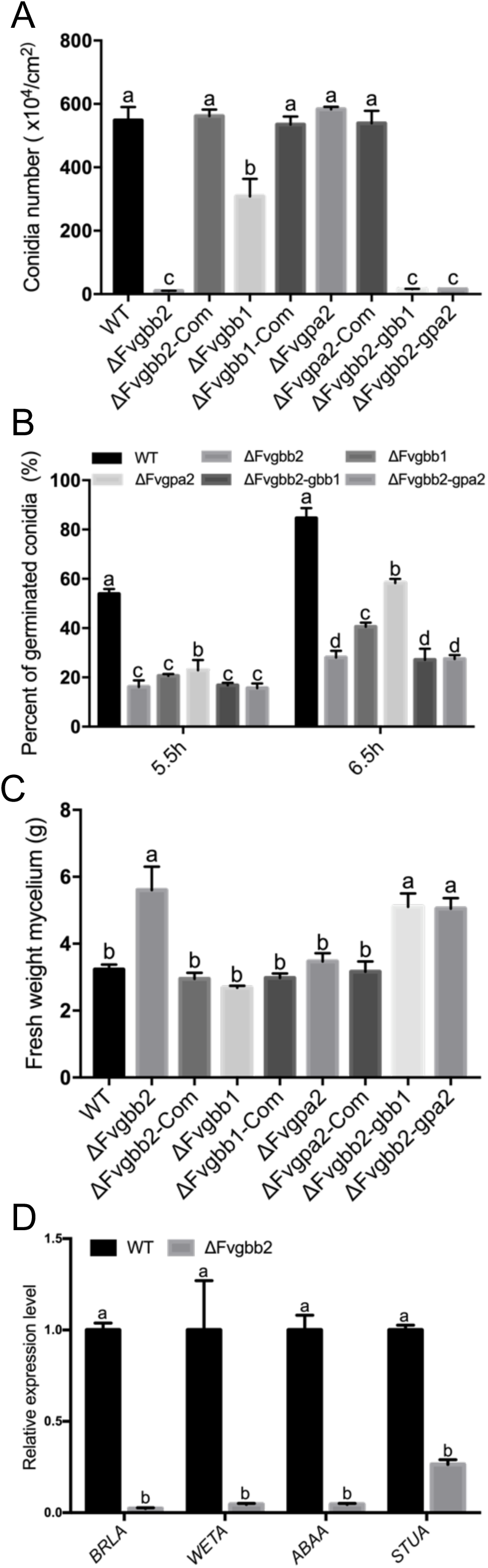
Impacts of gene deletion (ΔFvgbb2, ΔFvgbb1, ΔFvgpa2, ΔFvgbb2-gbb1, ΔFvgbb2-gpa2) on conidiation and vegetative growth. (A) Conidia production was assayed after 8-day incubation on V8 agar plates. Error bars indicate standard deviation from three replicates. The letters indicate statistically significant differences analyzed by Ordinary One-way ANOVA Fisher’s LSD test (p < 0.05). (B) WT, ΔFvgbb2, ΔFvgbb1, ΔFvgpa2, ΔFvgbb2-gbb1, ΔFvgbb2-gpa2 strains were suspended in 0.2xPDB liquid medium. Conidia showing germination were counted after 5.5 h and 6.5 h with gentle shaking. Different letters indicate significant difference according to a Two-Way ANOVA Fisher’s LSD test (P < 0.05). (C) Conidia (10^6^) in each strain was inoculated into 100 ml YEPD liquid medium, and images were taken after 7 days of shaking at 150 rpm. Fresh mycelia were harvested by filtering through Miracloth. Three replicates were performed for each strain. (D) Transcriptional level differences in conidia regulation genes in WT versus ΔFvgbb2 strain when cultured in myro liquid medium. Three replicates were performed for each sample.

### FvGBB2, FvGBB1 and FvGPA2 play important roles in secondary metabolism

Fumonisin B1 (FB1) is the most important mycotoxin produced by *F. verticillioides* in maize (Marin et al., 2013). Here, we assayed FB1 production in ΔFvgbb2, ΔFvgbb1, and ΔFvgpa2 in autoclaved kernels and surface-sterilized kernels (Silver Queen hybrid, Burpee Seeds) after 8 days of incubation (Fig. S4A and S4B). Interestingly, we noticed differences in pigment production (Fig. 4A and B). Additionally, FB1 production levels were dramatically lower in ΔFvgbb2, ΔFvgbb1 and ΔFvgpa2 mutants when compared to WT and complemented strains (Fig 4C and D). To verify the role of FvGbb2 in *F. verticillioides* secondary metabolism, we used qPCR to test transcription of 15 PKS genes. Total RNA samples were extracted from mycelia grown in myro broth, in which we noticed intense carmine red pigment production after 7 days of culturing with 150 rpm shaking (Fig. S4C). Consistently, all tested PKS genes showed significantly altered expressions in ΔFvgbb2. Notably, *PKS11* (*FUM1*), which is responsible for the first step of FB1 biosynthesis, was significantly downregulated in ΔFvgbb2 and ΔFvgbb1 (Table S1). Taken together, FvGbb2 is positively associated with FB1 production and necessary for maintaining proper regulation of other pigment productions.

**Fig. 4.**
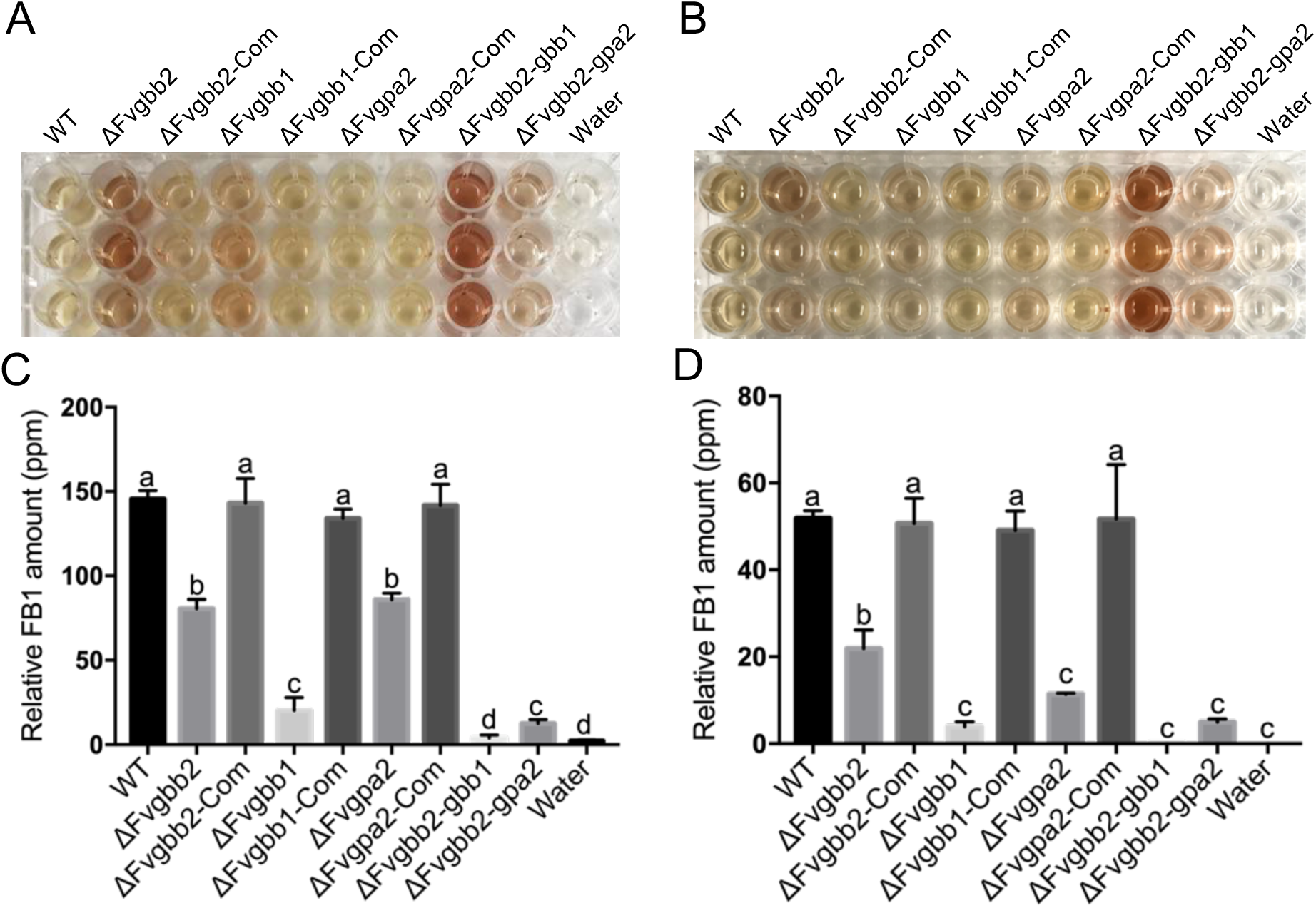
Role of *FvGBB2*, *FvGBB1*, *FvGPA2* on fumonisin B1 (FB1) and pigment productions. Pigment production observed in WT and deletion mutant strains grown on (A) autoclaved kernels and (B) surface sterilized kernels for 8 days. Samples were extracted overnight with 50 % acetonitrile (5 ml) and retrieved in a 96-well plate for the photograph. Three replicates were conducted for each strain. FB1 production in WT, mutants and complemented strains grown on (C) autoclaved kernels and (D) surface sterilized kernels for 8 days at room temperature. FB1 levels were normalized with fungal ergosterol levels.

### The subcellular localization of FvGbb2, FvGbb1, and FvGpa2

To study the subcellular localization of FvGbb2, we fused GFP to the C-terminus of FvGbb2 with its native promoter (Fig. S5A). We observed that strong FvGbb2-GFP signal widely distributed in *F. verticillioides* cells grown on both 0.2xPDA and myro agar medium (Fig. S5B∼D). To further uncover the relationship between FvGbb2 and G protein subunits, we examined colocalizations of FvGbb2 with FvGbb1 and FvGpa2. Briefly, mCherry-FvGbb1 and FvGpa2-mCherry were independently transformed into the FvGbb2-GFP strain. Unfortunately, we were not able to determine mCherry signals of FvGbb1 and FvGpa2 in growing hyphae tips (18 h) at room temperature due to very low signal intensity (data not shown). However, FvGbb1 and FvGpa2 mCherry signals from 0- and 10-h post-inoculation samples were clearly visible in the plasma membrane and vacuoles (Fig.5A and B). Significantly, our observations showed that GFP-FvGbb2 does not colocalize with mCherry-FvGbb1 and FvGpa2-mCherry, suggesting that FvGbb2 does not physically interact with FvGbb1 and FvGpa2 in selected time points in *F. verticillioides*.

**Fig. 5.**
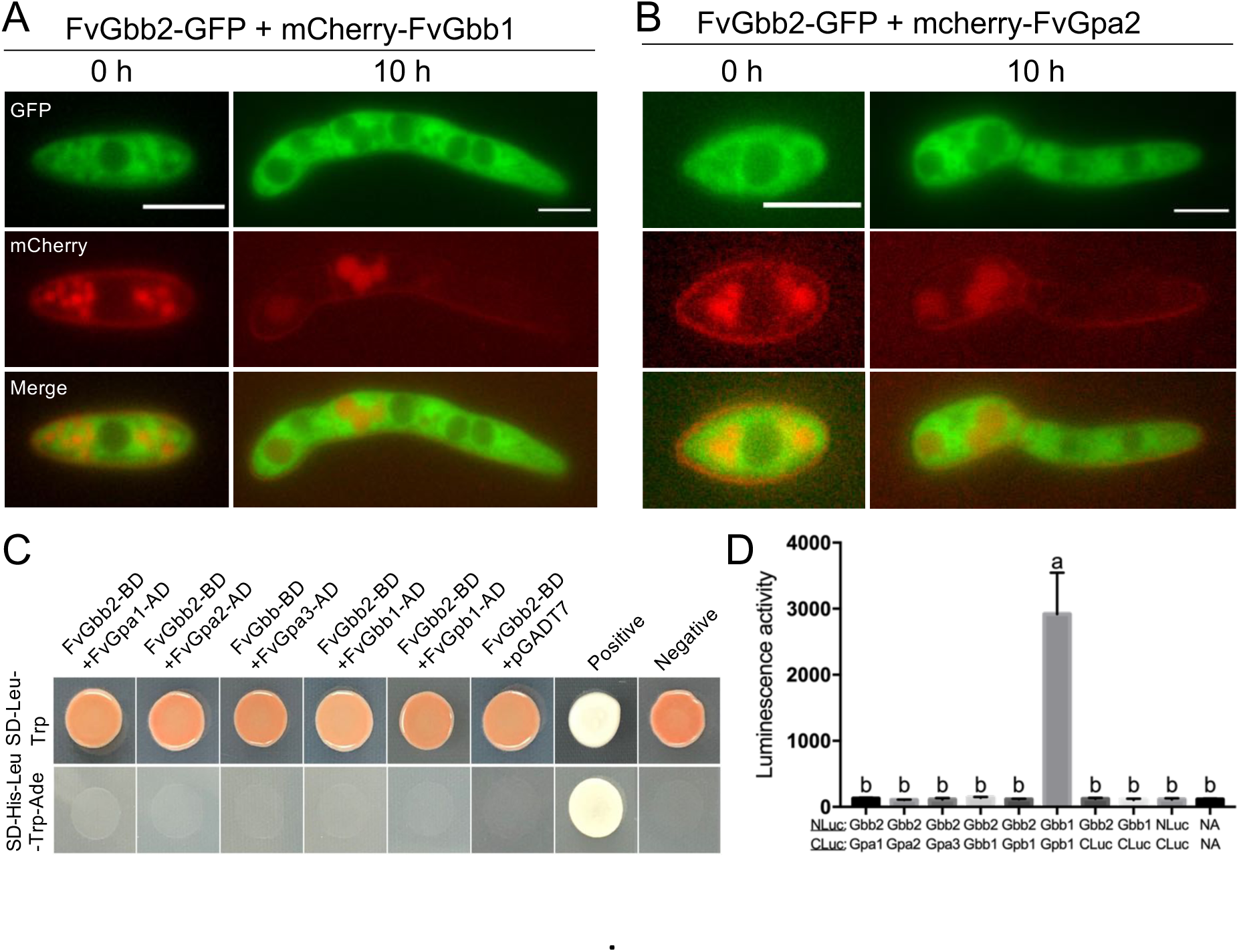
FvGbb2 localization and protein-protein interaction assays with FvGbb1 and FvGpa2 in *F. verticillioides*. Colocalization of FvGbb2-GFP and mCherry-FvGbb1 **(A)** and FvGbb2-GFP and FvGpa1-mCherry (B) are shown. We inoculated conidia (1×10^6^) on 0.2xPDA agar plates and cultivated at room temperature and took images at 0 h and 10 h post inoculation. Images were processed by ImageJ. Bar = 5μm. (C) Testing physical interaction between FvGbb2 and canonical heterotrimeric G protein components by yeast two-hybrid assay. FvGbb2 was co-introduced with Gα (FvGpa1, FvGpa2, and FvGpa3), Gβ (FvGbb1) and G*γ* (FvGpb1) into AH109 strain. Yeast transformants were grown on SD–His–Leu–Trp–Ade plates amended with 3 mm 3-AT. A pair of plasmids pGBKT7–53 and pGADT7-T was used as a positive control. A pair of plasmids pGBKT7–Lam and pGADT7-T was used a negative control. Images were taken after incubating at 30°C for 3 days. (D) Yeast two-hybrid assays were further verified by split luciferase complementation assays. Luminescence activity was acquired from three replicates and shown as relative light units (RLUs). A pair of FvGbb1-NLuc and FvGpb1-CLuc was used as a positive control. Multiple negative controls (FvGbb2-CLuc, FvGbb1-CLuc, NLuc-CLuc, no vector (NA)-NA) were also included in the study.

### *The Gβ-like protein FvGbb2 does not physically interact with* F. verticillioides *canonical heterotrimeric G protein components*

Our localization study led us to ask the relationship between FvGbb2 and canonical heterotrimeric G protein subunits. We used the yeast two-hybrid system to test the interaction between these proteins *in vivo*. Results showed that FvGbb2 does not physically interact with heterotrimeric G protein subunits (FvGpa1, FvGpa2, FvGpa3, FvGbb1, FvGpb1) in yeast cells (Fig. 5C). To further verify these results and to circumvent the limitations known in yeast two-hybrid system, we performed *in vivo* split luciferase complementation assay. However, we did not observe luciferase activity in all samples tested, indicating that FvGbb2 and G protein subunits (FvGpa1, FvGpa2, FvGpa3, FvGbb1, and FvGpb1) do not interact in *F. verticillioides* (Fig. 5D). FvGbb1 (Gβ subunit) and FvGpb1 (G*γ* subunit) were used as a positive control in split luciferase complementation assay.

### FvGbb2 is important for corn seedling virulence

Our previous study demonstrated that Gβ subunit is dispensable for the stalk rot virulence in *F. verticillioides* (Sagaram and Shim, 2007). FvGbb1 was fully capable of causing stalk rot disease when inoculated into maize B73 inbred stalks. To further investigate the function of Gβ-like protein on pathogenicity, we performed seedling rot assay with ΔFvgbb2, ΔFvgbb1, ΔFvgpa2, ΔFvgbb2-gbb1, ΔFvgbb2-gpa2 strains along with WT and complemented strains. The single mutant ΔFvgbb2 exhibited reduced virulence, but double mutants ΔFvgbb2-gbb1 and ΔFvgbb2-gpa2 were far less virulent when compared to the WT progenitor (Fig. 6A and 6B). These results suggest that FvGbb1, while physically not interacting with FvGbb2, plays a supplementary role in *F. verticillioides* seedling rot virulence.

**Fig. 6.**
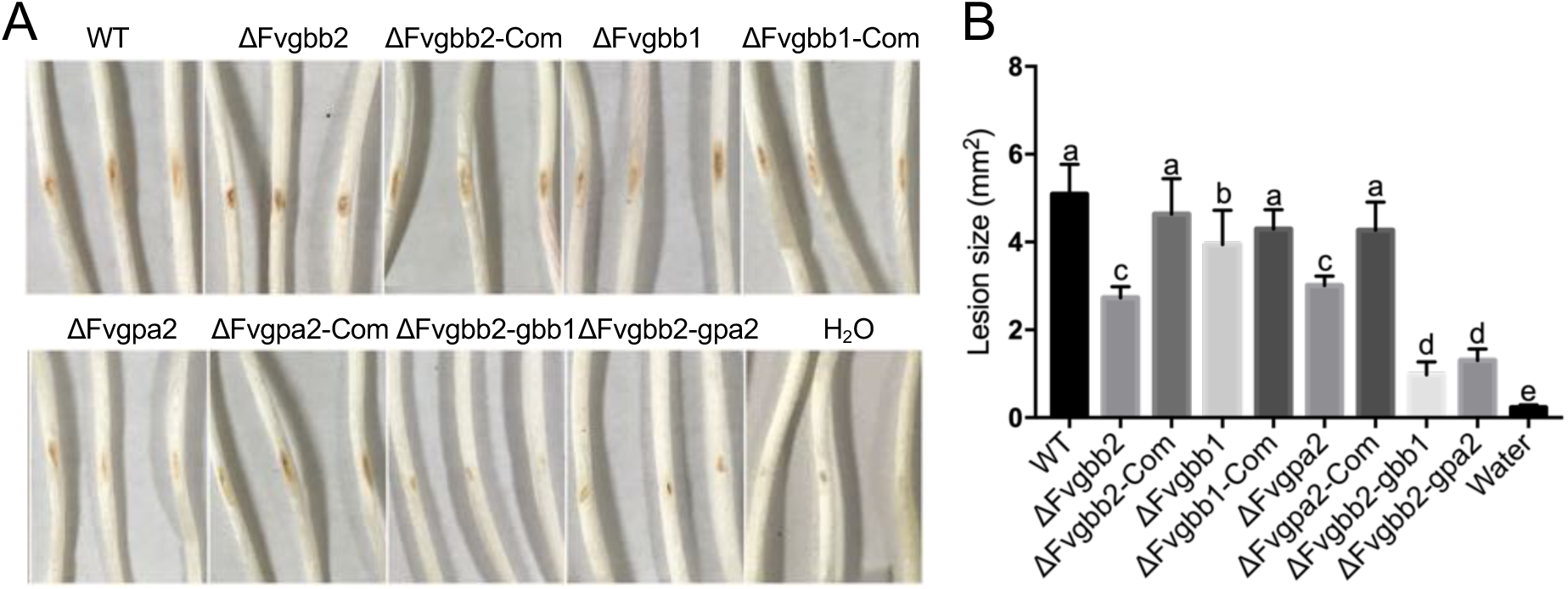
Attenuated virulence of Fvgbb2 mutant strain in the corn seedling rot assay. (A) Seedlings were grown in the dark room and inoculated with fungal spore suspension from each strain. The images were taken after a one-week incubation. (B) The lesion size of seedling rot was examined and measured by image J. At least three biological and three technical replicates were assayed for each strain.

### FvGbb2 is associated with carbon source utilization and stress response

Yeast RACK1 homolog ScAsc1 is positively associated with glucose sensing (Zeller et al., 2007). Here, we cultured mutant strains on different carbon sources to test whether FvGbb2 played a similar role in *F. verticillioides*. In addition to dextrose, we tested sucrose, fructose and xylose as the sole carbon source in Czapek-Dox agar plates (Fig. 7A and C). While ΔFvgbb2 showed general vegetative growth defect, we learned that the negative impact on mycelial growth was more severe in dextrose when compared to sucrose, fructose and xylose. This result was in agreement with the study in *S. cerevisiae*. Additionally, ΔFvgbb2 exhibited a similar pattern related to growth in fructose but to a lesser extent in contrast to xylose medium.

**Fig. 7.**
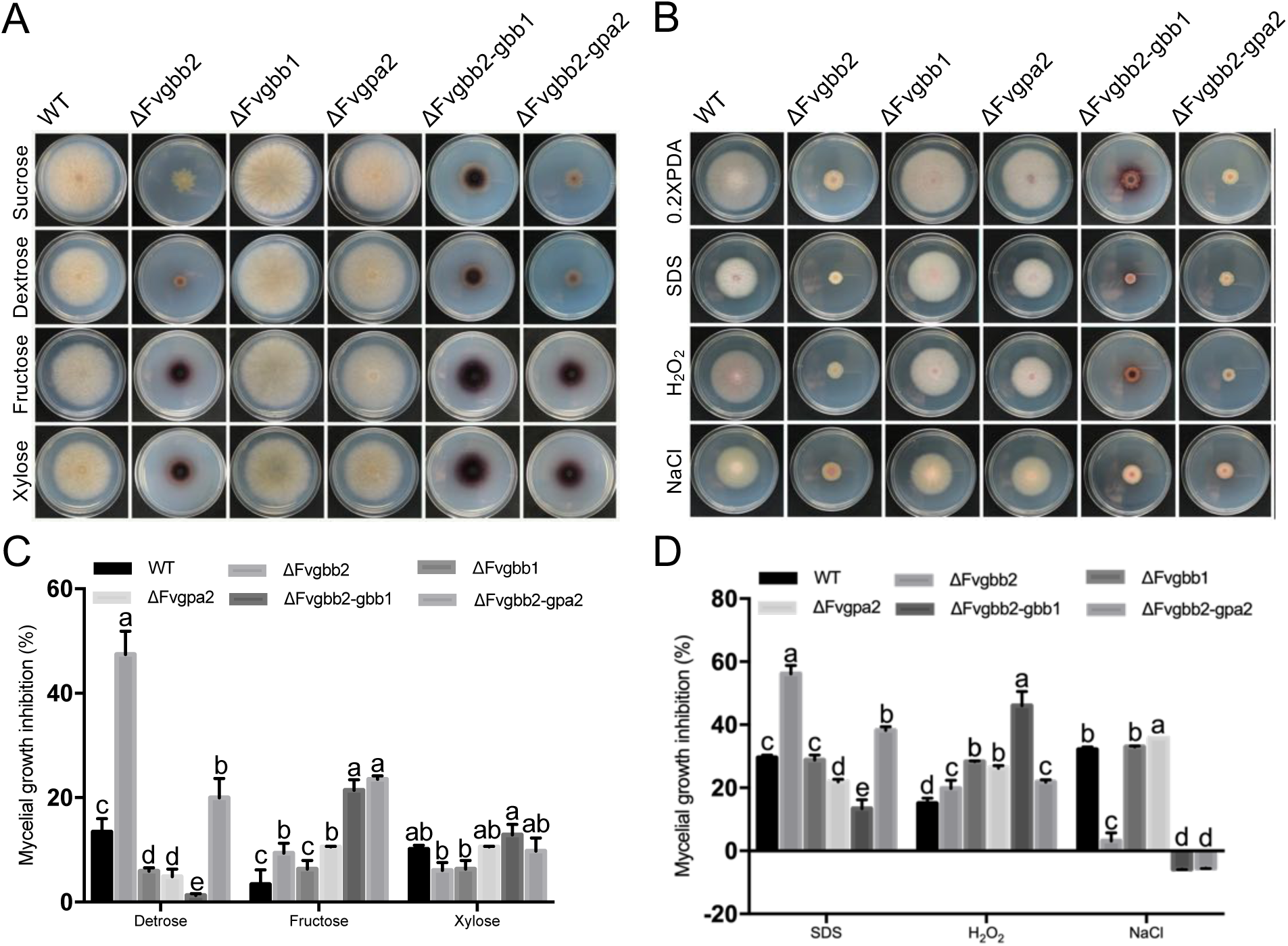
Influences on deletion of *FvGBB2*, *FvGBB1*, *FvGPA2* to carbon utilization and stress agents. (A) Strains were cultivated on modified Czapek-Dox agar with different carbon source, including sucrose, dextrose, fructose, and xylose, and incubated at room temperature for 8 days. (B) Radial growth of WT and deletion mutants with various stress agents amended in 0.2xPDA agar plates incubated for 8 days at room temperature. (C) The growth inhibition rate was analyzed by comparing the growth on Czapek-Dox agar with sucrose against other carbon amendments. The mycelial growth inhibition rate (%) was measured by ((sucrose growth diameter - designated carbon diameter) / sucrose growth diameter) x 100. Bar indicates standard deviation of three replicates. (D) The inhibition rate (%) of strains of stress responses was measured by ((0.2xPDA growth diameter - designated stress growth diameter) / 0.2xPDA growth diameter) x 100. Bar indicates standard deviation of three replicates.

**Fig. 8.**
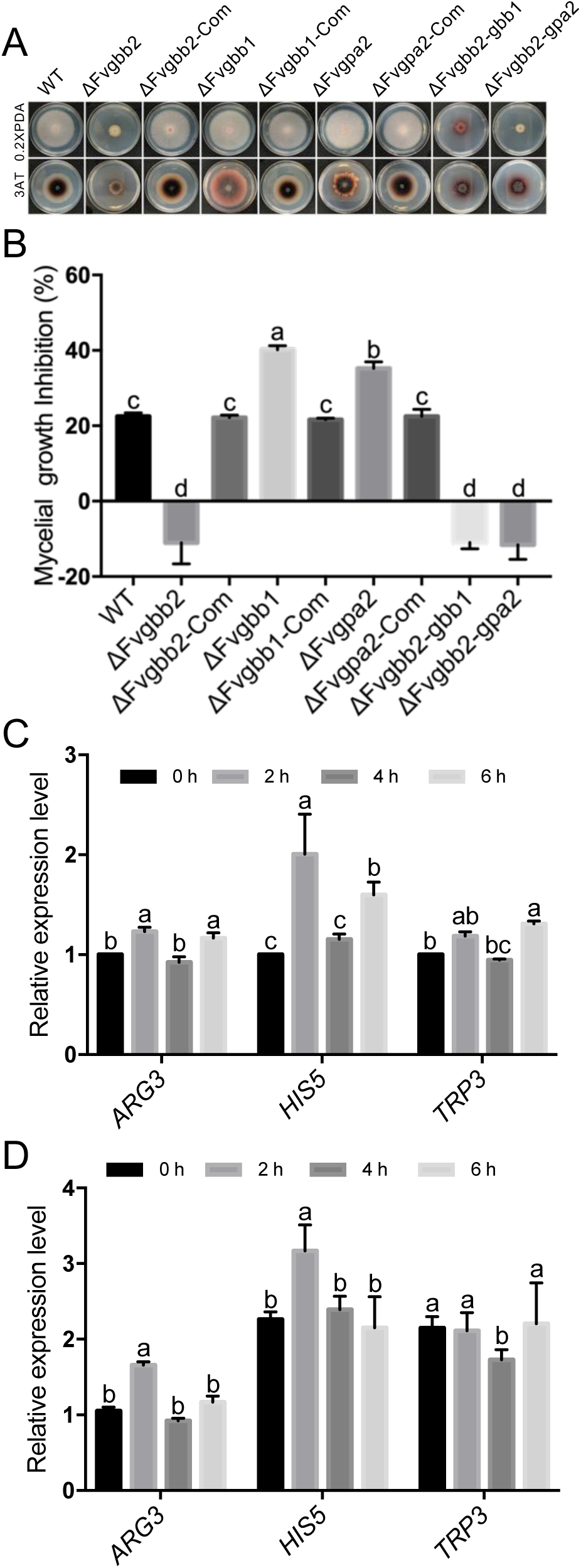
Negative impacts of FvGbb2 on amino acid starvation. (A) Strains were cultured on 0.2xPDA without and with 3mM 3-AT for 8 days at room temperature. (B) Mycelial growth inhibitions of WT and mutant strains were compared between 0.2xPDA without and with 3mM 3-AT. (C) Transcription levels of *ARG3*, *HIS5* and *TRP3* in WT were analyzed by qPCR after treatment with 3mM 3-AT for 0h, 2h, 4h, 6h by qPCR. (D) Relative expression levels of three GAAC related genes were studied by qPCR in ΔFvgbb2 after treatment with 3mM 3-AT for 0h, 2h, 4h, 6h.

Moreover, in *S. cerevisiae*, MAPK Slt2 phosphorylation level was enhanced *Scasc1* mutant (Chasse et al., 2006). Thus, we hypothesized that FvGbb2 is associated with stress responses in *F. verticillioides*. As shown in Fig. 7B and D, ΔFvgbb2 exhibited increased sensitivity to SDS (cell wall stress) and H_2_O_2_ (oxidative stress) but not to NaCl (salt stress). When relatively compared, ΔFvgbb2 grew better in NaCl than with SDS or H_2_O_2_ in the medium. This result indicates that FvGbb2 is positively associated with cell wall stress response but negatively with salt tolerance. To test whether FvGbb2 regulates stress response via interacting with MAP kinase pathways, we performed yeast two-hybrid and split luciferase complementation assays to test the interaction between FvGbb2 with three cell wall integrity MAPK kinase cascade proteins (FvSlt2, FvMkk1/2, FvBck1) (Fig. S6A and S6B). However, these studies did not reveal direct physical interactions.

### FvGbb2 is negatively involved in amino-acid starvation

In response to amino acid starvation, *S. cerevisiae* cells stimulate the expression of a number of amino acid metabolism genes, which is known as the general amino acid control (GAAC) response (Wek et al., 2004). Here, we performed a growth assay in the medium containing 3 mM 3-amino-l,2,4-triazole (3-AT) to determine the possibility of FvGbb2 playing a role in GAAC response. As shown in Fig 9A and 9B, ΔFvgbb2, ΔFvgbb2-gbb1 and ΔFvgbb2-gpa2 stains all showed similar resistance to 3-AT treatment. To investigate whether the growth suppression in ΔFvgbb2 is due to the increase in transcription of amino acid biosynthesis genes, we tested the expression of three genes associated with GAAC response. Consistently, *HIS5* and *TRP3* genes were up-regulated in the ΔFvgbb2. Notably, *HIS5* and *TRP3* gene transcription had doubled under a non-starving condition (0h with no 3-AT) in ΔFvgbb2 compared to WT. But we did not observe *ARG3* gene expression increase with 3-AT treatment in ΔFvgbb2 in contrast to WT at selected time points. These results suggest that FvGbb2 plays an important role in suppressing GAAC response in *F. verticillioides*, which is similar to *S. cerevisiae*.

## Discussion

Our previous study characterized *F. verticillioides* Gβ protein FvGbb1, a key component of the canonical heterotrimeric G protein complex, that demonstrated an important role for FB1 biosynthesis. However, despite the roles Gβ subunit is known to play in the heterotrimeric G protein complex, we were quite baffled by the fact that FvGbb1 null mutation had little impact on vegetative growth, sexual and asexual reproduction, and pathogenicity in *F. verticillioides* (Sagaram and Shim, 2007). Particularly, Gβ protein subunits in other fungal species are known to play some critical roles. For instance, mutational inactivation studies in *M. oryzae*, *C. neoformans* and *N. crassa* indicated that Gβ proteins were critically associated with sexual mating while were responsible for virulence in *M. oryzae* and *F. graminearum* (Rosen et al., 1999; Wang et al., 2000; Nishimura et al., 2003; Yu et al., 2008). Namely, *M. oryzae* Mgb1 deletion mutant showed defects in appressorium formation and thus failed to penetrate host plant leaves. It was also observed that the mutation led to cAMP level reductions in *M. oryzae* cells and that exogenous application of cAMP failed to restore appressorium functions. Moreover, we must recognize that Gβ protein is one of the well-characterized WD40-repeat proteins, and numerous published reports demonstrate important physiological and genetics roles WD40-repeat proteins play in filamentous fungi. For instance, fungal striatin-like proteins are important WD40-repeat proteins known to function as a scaffolding protein required for fungal anastomosis, sexual development and virulence in *N. crassa*, *Sordaria macrospora*, and *Fusarium* species (Shim et al., 2006; Bloemendal et al., 2012; Dettmann et al., 2013)

Based on our experimental data and literature, we hypothesized the presence of a Gβ-like protein that can supplement or perhaps substitute canonical Gβ subunit in *F. verticillioides*. In this study, we identified FvGbb2, which shares structural similarity with FvGbb1 with seven WD40 repeat domains. As we mentioned earlier, since Gβ protein FvGbb1 does not play a critical role in *F. verticillioides* growth and virulence (Sagaram and Shim, 2007), we tested whether FvGbb2 serve these functions. The Gβ subunit is a well-recognized regulator of asexual development and secondary metabolism in fungi. As shown in this study, the deletion mutant of *FvGBB2* exhibited severe defects in growth and conidiation in solid medium. Our qPCR results also revealed that transcription of conidia-related genes including *BRLA* are highly suppressed in tested media. In *Aspergillus* species, transcriptional and metabolic studies indicated that *BRLA* is associated with regulating secondary metabolisms in addition to as a well-known master element in asexual development (Adams and Yu, 1998; Lind et al., 2018). This result raised a question whether secondary metabolites such as mycotoxins and pigments in *F. verticillioides* also follow the same pattern of regulation as in *Aspergillus*. Additionally, earlier studies also demonstrated that Gβ-like proteins positively regulate mycotoxin production in *F. graminearum* and *A. fumigatus*. Similarly, we found that the deletion of *FvGBB2* led to a reduction in FB1 production and enhanced the production of pigments in maize (Fig 4A), myro liquid medium (Fig 4C), and 0.2xPDB (data not shown). However, it is interesting to note that the pigment biosynthesis was not altered in *F. graminearum* when *Δ*Fgasc1 was cultured in PDB (Tang et al., 2018). When we performed expression analyses of 15 PKSs genes, we learned that most PKSs genes are significantly downregulated, including *PKS11* (*FUM1*). And we also saw overexpression of select PKS genes which may explain why we observe a high level of pigment production in ΔFvgbb2 cultures. This is contradictory to results observed in *A. nidulans*. *i.e.* Gβ-like protein inactivation did not alter mycotoxin sterigmatocystin (ST) biosynthesis *stcU* gene expression in *A. nidulans* (Kong et al., 2013). We concluded that FvGbb2 regulates transcriptional activities of key conidia-related genes and impacts fungal secondary metabolism and asexual reproduction, as is the case with canonical heterotrimeric Gβ subunit.

Previous studies in *Candida albicans* and *A. fumigatus* demonstrated that the deletion of RACK1 leads to a dramatic reduction of virulence when tested in animal models (Liu et al., 2010; Cai et al., 2015). Consistent with these results, ΔFvgbb2 exhibited approximately 50% reduction in virulence when tested for maize stalk rot. The virulence deficiency likely is due to the impaired vegetative growth on seedlings. However, we did not rule out the possibility that ΔFvgbb2 virulence deficiency is also due to the impaired response against stress factors. In *S. cerevisiae* and *M. oryzae*, the RACK1 deletion mutants showed higher sensitivity when exposed to cell wall stress agents (Rachfall et al., 2013; Li et al., 2017). Notably, *A. thaliana* RACK1 homolog has been shown to interact with three tiers of the MAPK cascade components, while in *M. oryzae* the protein interacts with MoBck1 (Cheng et al., 2015; Li et al., 2017). These MAPK pathways are known to be important signal transduction pathways in eukaryotes (Hamel et al., 2012). Similar to *S. cerevisiae* and *M. oryzae*, ΔFvgbb2 exhibited sensitivity towards SDS stress, but we found no experimental evidence that FvGbb2 physically interacts with MAPK kinase cascade components FvBck1, FvMkk1/2, and FvSlt2. However, we are further investigating this due to the possibility that FvGbb2-MAPK interactions may occur in a specific, yet-to-be determined condition. On another note, the RACK1 homolog in *S. pombe* Cpc2 was positively involved in the cytoplasmic catalase production, which in turn mediates the detoxification of hydrogen peroxides (Nunez et al., 2009). In line with this previous study, ΔFvgbb2 exhibited more sensitivity to oxidative stress agents. Moreover, a prior study demonstrated that rice OsRACK1A has negative impacts on salt tolerance and interacts physically with salt stress-related proteins (Zhang et al., 2018a). Our analysis also revealed that ΔFvgbb2 mutant strain showed a higher level of tolerance against NaCl treatment. In *A. fumigatus,* the mutation in *AfcpcB* gene showed no such effect (Cai et al., 2015).

These stress responses observed in *F. verticillioides* could be explained by the proteome and transcriptome studies in *S. cerevisiae* since the mutation in RACK1 homolog Asc1 impacted the expression of various regulatory response components including those in MAPK kinase signal pathway (Rachfall et al., 2013).

RACK1 shares structural similarity, and therefore is hypothesized to function as a Gβ subunit and physically interact with Gα subunits. In yeast, the interaction with one of the two Gα subunits, Gpa2, was documented while RACK1 homolog in *F. graminearum* Gib2 was shown to interact with one of three Gα proteins, Gpa1, and two G*γ* subunits (Palmer et al., 2006; Zeller et al., 2007). Interestingly, two studies in *A. thaliana* were not in agreement when characterizing the relationship between RACK1 homolog and canonical G protein components. While one previous report in *A. thaliana* using yeast two-hybrid and *in vivo* Co-IP assay failed to detect interactions between RACK1 and G proteins, the other research group showed that scaffolding protein RACK1 interacted with Gβ protein in *A. thaliana* validated by bimolecular fluorescence, split firefly luciferase complementation and co-immunoprecipitation (Guo et al., 2009; Cheng et al., 2015). In our study, direct interaction between FvGbb2 with canonical heterotrimeric G proteins could not be established by yeast two-hybrid and split luciferase complementation assays. We also could not observe colocalization of FvGbb2-GFP with mCherry-FvGbb1 and FvGpa2-mCheery. However, ΔFvgbb2-gbb1 and ΔFvgbb2-gpa2 showed more severe defects in FB1 production and pathogenicity when compared to single mutant strains. We propose that although Gβ protein FvGbb1 and Gβ-like protein FvGbb2 share similarity, their localizations and functions are distinct.

While FvGbb1 is expressed mostly in the vacuole in conidia and early elongation stage, FvGbb2 is highly expressed in the cytoplasm constitutively. We observed diverse roles FvGbb2 plays in *F. verticillioides*, and this may be explained by how FvGbb2-GFP signal is distributed broadly in the cytoplasm. Thus, it would be reasonable to hypothesize that FvGbb2 interacts with diverse partners in the cell concurrently or sequentially, and in turn regulates differential gene expression and translation of downstream proteins associated with vegetative growth, virulence, mycotoxin biosynthesis, carbon utilization and stress response in *F. verticillioides*.

## Experimental procedures

### Fungal strains and growth study

The *F. verticillioides* strain M3125 was used as a wild-type strain in this study. All strains in this study were cultivated on V8, 0.2xPDA and myro agar plates as described previously (Yan et al., 2019). For spore gemination assay, equal amounts of newly harvested microconidia grown on V8 agar plates for 8 days were cultivated in 0.2x potato dextrose broth (PDB) (Sigma-Aldrich) for 5.5 h and 6.5 h with gentle shaking. For stress assay, strains with 4 μl of 1 × 10^6^ conidial suspension were grown on 0.2xPDA agar plates with various stressors including 0.01% SDS, 2 mM H_2_O_2_, and 0.75 M NaCl. For carbon utilization assay, Czapek-Dox agar with four different carbon sources in Czapek-Dox agar as follows: sucrose (30g/L), dextrose (10g/L), fructose (10g/L) and xylose (10g/L) (Yan et al., 2019). Fungal growth diameters were determined after 8 days of incubation at room temperature. Perithecia formation was conducted by applying the same amount of spores obtained from V8 agar plates and spread to the strain 3120 on carrot plates following the previous method (Sagaram and Shim, 2007). For the amino acid starvation assay, 0.2xPDA with 3mM 3-amino-l,2,4-triazole (3AT) were used for testing the growth impact. We inoculated 0.5 ml of spores (10^6^) WT and mutants into 50 ml YEPD at constant shaking for 22 h. Following shaking, 3AT was added to bring the final concentration to 3mM. Time point samples were collected after continuous gently shaking at 2h, 4h and 6h. All experiments had at least three replicates.

### Gene deletion and complementation

Knockout mutants of **Δ**Fvgbb2, **Δ**Fvgbb1, **Δ**Fvgpa2 were generated in *F. verticillioides* M3125 strain via homologous recombination using split-marker approach. A hygromycin B phosphotransferase gene (*HPH*) designated as *PH* (929bp) and *HP* (765bp) or a geneticin resistance gene (*GEN*) designated as *GE* (1183 bp) and *EN* (1021 bp) were used with joint-PCR to fuse with gene left and right flanking regions. For generating double knockout mutants, *FvGPA2* and *FvGBB1* genes were replaced with *GEN* gene in **Δ**Fvgbb2 background using the same strategy as described above, respectively. For complementation, we used the native promoter followed by the gene and terminator cotransforming with either pBSG (*GEN*) or pBP15 (*HPH*) plasmids to the single mutant background. All knockout constructs and complementation fragments in this study were amplified using Phusion Flash High-Fidelity PCR Master Mix (Thermo Scientific™: F548L) following the manufacturer’s instructions. All transformants were screened by PCR using Phire Plant Direct PCR Kit (Thermo Scientific™: F130WH or F122S) and Taq DNA polymerase (NEB: M0267S) to identify putative mutant strains. qPCR was used for further confirmations. All primers used in this study were shown in Table S2.

### Construction of fluorescent strains

For constructing FvGbb2-GFP plasmid, Fv*GBB2* fragment (2818 bp) was amplified from the genomic DNA of *F. verticillioides* with the primers FvGbb2-GFP-F/R using Q5® High-Fidelity DNA Polymerase (NEB: M0491S). The DNA fragment was introduced to the pKNTG plasmid KpnI and HindIII sites via In-Fusion-HD cloning kit (Takara Bio USA, Inc). We sequenced plasmids and transformed into the ΔFvgbb2 and WT strain resulting in ΔFvgbb2-Gbb2-GFP and FvGbb2-GFP strain. To construct the mCherry-FvGbb1 plasmid, FvGbb1 native promoter was amplified from genomic DNA. cDNA was then used to amplify FvGbb1 coding sequence and 3’ UTR. pKNT-mCherry was used for mCherry fragment amplification. These three PCR products were cloned into KpnI and BamHI sites of pKNT. Based on suggestions in Gα protein florescent strain constructions, mCherry was tagged in the amino acid 114 site to construct FvGpa2-mCherry plasmid (Eaton et al., 2012; Ramanujam et al., 2013). The mCherry-FvGbb1 and FvGpa2-mCherry were cotransformed with hygromycin fragment into FvGbb2-GFP strain, respectively. All DNA fragments used in this study were purified using GeneJET Gel Extraction Kit (Thermo Scientific™). Plasmids in this study were isolated using GeneJET Plasmid Maxiprep Kit (Thermo Scientific™).

### Yeast two-hybrid and split luciferase complementation assay

The coding sequence of *FvGPA1* (FVEG_06962), *FvGPA2* (FVEG_04170), *FvGPA3* (FVEG_02792), *FvGPB1*(FVEG_05349), *FvGBB1* (FVEG_10291)*, FvMKK1/2* (FVEG_05280), *FvSLT2* (FVEG_03043), and *FvBCK1*(FVEG_05000) were used to construct plasmids for yeast two-hybrid assay. These were amplified from *F. verticillioides* strain M3125 cDNA using Q5® High-Fidelity DNA Polymerase and inserted in pGADT7 (Clontech, Mountain View, CA, USA) as prey vectors. The full length of the coding region of FvGbb2 was cloned into pGBKT7 as a bait vector. The resulting plasmids were verified by sequencing. The pair of yeast two-hybrid plasmids were transformed into yeast strain AH109 following the instruction. We added 5 μl of transformants (10^7^/mL) on SD/–Leu/–Trp (Clontech, Mountain View, CA, USA) and SD/–Ade/– His/–Leu/–Trp (3 mM 3-AT) agar plates.

Split luciferase complementation assay plasmids in this study were obtained as described previously (Kim et al., 2012). The coding region of each gene was amplified from WT strain cDNA by Q5® High-Fidelity DNA Polymerase and inserted into pFNLucG or pFCLucH via In-Fusion® HD Cloning (Clontech, Mountain View, CA, USA). The resulting constructs were validated by sequencing. Transformation, selection and luciferin assay were followed by a previous description (Zhang et al., 2018b).

### Fumonisin B1 and pathogenicity assays

For determining the FB1 production, cracked corns (2 g) were put in 20 mL scintillation vials (VWR) and rehydrated with 1 mL sterilized water overnight followed by autoclaving. Fungal spore solutions (200 μL, 10^6^/mL) were inoculated in each vial and cultivated at room temperature for eight days. Additionally, four silver queen kernels were sterilized using the previous description, but 10% bleach was used instead of 6% sodium hypochlorite (Christensen et al., 2012). Sterilized living seeds were placed on the sterilized 90 mm Whatman filter paper and created a wound on the endosperm area by a scalpel. Fungal spore solution (100 μL, 10^6^/mL) were inoculated in each vial and grown at room temperature for eight days. FB1 and ergosterol extraction methods were described previously (Christensen et al., 2012) with a minor modification. In this study, we used 5 mL of 50% acetonitrile for FB1 extraction while 5ml of chloroform: methanol (2:1, v/v) for ergosterol extraction. HPLC analyses of FB1 and ergosterol were performed as described (Shim and Woloshuk, 1999). FB1 levels were then normalized to ergosterol contents. These experiments were carried out with three biological replicates. Stalk virulence was conducted on silver queen hybrid seedling (Burpee) as previously described on corn seedling with minor modifications (Kim et al., 2018). Spores solution (5 μL, 10^7^/mL) were collected from V8 plates. The seedlings were collected and imaged after a one-week growth in the dark room.

### RNA extraction and relative gene expression study

For a 20 h qPCR study assay, a 2 mL conidia suspension (10^6^ conidia/mL) was inoculated in 100 mL YEPD liquid medium and incubated at 150 rpm agitation for 3 days at room temperature. For myro assay, a 200 μl conidia solution (10^6^ conidia/ml) was inoculated in 100 ml YEPD liquid medium for 3 days. Subsequently, mycelium was harvested through Miracloth (EMD Millipore) and weighed 0.3g to 100ml myro liquid medium. Samples were collected after 7 days incubating at 150 rpm. Three replicates were conducted for each strain. Total RNA was isolated using Qiagen RNA Plant Mini kit following the kit instructions. For qPCR, cDNA was synthesized using the Verso cDNA synthesis kit (Thermo Fisher Scientific). qPCR analyses were conducted on Step One plus real-time PCR system using the DyNAmo ColorFlash SYBR Green qPCR Kit (Thermo Fisher Scientific). Relative expression levels of each gene were calculated using a 2^−ΔΔCT^ method and normalized with *F. verticillioides β*-tubulin-encoding gene (FVEG_04081). All qPCR assays were performed with three replicates. PKs primers were from a previous study (Ortiz and Shim, 2013).

### Microscopy

For hyphal branching imaging, minor modifications were made from a previous study (Schultzhaus et al., 2015). Briefly, strains were cultivated on 0.2xPDA agar plates for three days. A block of agar was cut and put on a glass slide. Sterilized water (10 μL) water was added followed by a coverslip. The sample was incubated at room temperature for 20 mins and examined by a microscope (Olympus BX60). For GFP and mCherry assay, we used the same method but indicated time conidia and hypha were used to image instead. Images were processed using ImageJ software (Schneider et al., 2012).

## Acknowledgements

This research was supported in part by the Agriculture and Food Research Initiative Competitive Grants Program Grant (2013-68004-20359) from the USDA National Institute of Food and Agriculture. We thank Dr. Brian Shaw, Ms. Blake Commer and Mr. Joe Vasselli in the Department of Plant Pathology & Microbiology, Texas A&M University for assisting in microscopy. We thank Dr. Wenhui Zheng (Fujian Agriculture & Forestry University) for providing pKNTG and pKNT-mCherry plasmids. We thank Dr. Meilian Chen (Fujian Agriculture & Forestry University) for her advice in the yeast two-hybrid experiment. The authors declare no conflict of interest.

## References

1. Adams, T.H., and Yu, J.H. (1998) Coordinate control of secondary metabolite production and asexual sporulation in *Aspergillus nidulans*. Curr Opin Microbiol 1: 674–677.

2. Blacutt, A.A., Gold, S.E., Voss, K.A., Gao, M., and Glenn, A.E. (2018) *Fusarium verticillioides*: Advancements in Understanding the Toxicity, Virulence, and Niche Adaptations of a Model Mycotoxigenic Pathogen of Maize. Phytopathology 108: 312–326.

3. Bloemendal, S., Bernhards, Y., Bartho, K., Dettmann, A., Voigt, O., Teichert, I. et al. (2012) A homologue of the human STRIPAK complex controls sexual development in fungi. Mol Microbiol 84: 310–323.

4. Cai, Z.-d., Chai, Y.-f., Zhang, C.-y., Qiao, W.-r., Sang, H., and Lu, L. (2015) The Gβ-like protein CpcB is required for hyphal growth, conidiophore morphology and pathogenicity in *Aspergillus fumigatus*. Fungal Genetics and Biology 81: 120–131.

5. Chasse, S.A., Flanary, P., Parnell, S.C., Hao, N., Cha, J.Y., Siderovski, D.P., and Dohlman, H.G. (2006) Genome-scale analysis reveals Sst2 as the principal regulator of mating pheromone signaling in the yeast *Saccharomyces cerevisiae*. Eukaryotic Cell 5: 330–346.

6. Cheng, Z., Li, J.-F., Niu, Y., Zhang, X.-C., Woody, O.Z., Xiong, Y. et al. (2015) Pathogen-secreted proteases activate a novel plant immune pathway. Nature 521: 213-+.

7. Christensen, S., Borrego, E., Shim, W.B., Isakeit, T., and Kolomiets, M. (2012) Quantification of fungal colonization, sporogenesis, and production of mycotoxins using kernel bioassays. J Vis Exp.

8. Coyle, S.M., Gilbert, W.V., and Doudna, J.A. (2009) Direct Link between RACK1 Function and Localization at the Ribosome In Vivo. Molecular and Cellular Biology 29: 1626–1634.

9. Dettmann, A., Heilig, Y., Ludwig, S., Schmitt, K., Illgen, J., Fleissner, A. et al. (2013) HAM-2 and HAM-3 are central for the assembly of the *Neurospora* STRIPAK complex at the nuclear envelope and regulate nuclear accumulation of the MAP kinase MAK-1 in a MAK-2-dependent manner. Mol Microbiol 90: 796–812.

10. Eaton, C.J., Cabrera, I.E., Servin, J.A., Wright, S.J., Cox, M.P., and Borkovich, K.A. (2012) The guanine nucleotide exchange factor RIC8 regulates conidial germination through Gα proteins in *Neurospora crassa*. PLoS One 7: e48026.

11. Guo, J., Wang, S., Wang, J., Huang, W.D., Liang, J., and Chen, J.G. (2009) Dissection of the relationship between RACK1 and heterotrimeric G-proteins in *Arabidopsis*. Plant Cell Physiol 50: 1681–1694.

12. Hamel, L.P., Nicole, M.C., Duplessis, S., and Ellis, B.E. (2012) Mitogen-activated protein kinase signaling in plant-interacting fungi: distinct messages from conserved messengers. Plant Cell 24: 1327–1351.

13. Hansen, F.T., Gardiner, D.M., Lysoe, E., Fuertes, P.R., Tudzynski, B., Wiemann, P. et al. (2015) An update to polyketide synthase and non-ribosomal synthetase genes and nomenclature in *Fusarium*. Fungal Genet Biol 75: 20–29.

14. He, Z., Zhang, H., Gao, S., Lercher, M.J., Chen, W.H., and Hu, S. (2016) Evolview v2: an online visualization and management tool for customized and annotated phylogenetic trees. Nucleic Acids Res 44: W236–241.

15. Keller, N.P. (2018) Fungal secondary metabolism: regulation, function and drug discovery. Nat Rev Microbiol.

16. Kim, H.K., Cho, E.J., Jo, S., Sung, B.R., Lee, S., and Yun, S.H. (2012) A split luciferase complementation assay for studying in vivo protein-protein interactions in filamentous ascomycetes. Curr Genet 58: 179–189.

17. Kim, M.S., Zhang, H., Yan, H., Yoon, B.J., and Shim, W.B. (2018) Characterizing co-expression networks underpinning maize stalk rot virulence in *Fusarium verticillioides* through computational subnetwork module analyses. Sci Rep 8: 8310.

18. Kong, Q., Wang, L., Liu, Z., Kwon, N.-J., Kim, S.C., and Yu, J.-H. (2013) Gβ-Like CpcB Plays a Crucial Role for Growth and Development of *Aspergillus nidulans* and *Aspergillus fumigatus*. Plos One 8.

19. Kumar, S., Stecher, G., and Tamura, K. (2016) MEGA7: Molecular Evolutionary Genetics Analysis Version 7.0 for Bigger Datasets. Mol Biol Evol 33: 1870–1874.

20. Larkin, M.A., Blackshields, G., Brown, N.P., Chenna, R., McGettigan, P.A., McWilliam, H. et al. (2007) Clustal W and Clustal X version 2.0. Bioinformatics 23: 2947–2948.

21. Letunic, I., and Bork, P. (2018) 20 years of the SMART protein domain annotation resource. Nucleic Acids Res 46: D493–D496.

22. Letunic, I., Doerks, T., and Bork, P. (2015) SMART: recent updates, new developments and status in 2015. Nucleic Acids Res 43: D257–260.

23. Li, G., Zhang, X., Tian, H., Choi, Y.E., Tao, W.A., and Xu, J.R. (2017) *MST50* is involved in multiple MAP kinase signaling pathways in *Magnaporthe oryzae*. Environ Microbiol 19: 1959–1974.

24. Li, L., Wright, S.J., Krystofova, S., Park, G., and Borkovich, K.A. (2007) Heterotrimeric G protein signaling in filamentous fungi. In Annual Review of Microbiology, pp. 423–452.

25. Lind, A.L., Lim, F.Y., Soukup, A.A., Keller, N.P., and Rokas, A. (2018) An LaeA- and BrlA-Dependent Cellular Network Governs Tissue-Specific Secondary Metabolism in the Human Pathogen Aspergillus fumigatus. mSphere 3.

26. Liu, X., Nie, X., Ding, Y., and Chen, J. (2010) Asc1, a WD-repeat protein, is required for hyphal development and virulence in *Candida albicans*. Acta Biochimica Et Biophysica Sinica 42: 793–800.

27. Ma, L.J., Geiser, D.M., Proctor, R.H., Rooney, A.P., O’Donnell, K., Trail, F. et al. (2013) *Fusarium* pathogenomics. Annu Rev Microbiol 67: 399–416.

28. Marin, S., Ramos, A.J., Cano-Sancho, G., and Sanchis, V. (2013) Mycotoxins: occurrence, toxicology, and exposure assessment. Food Chem Toxicol 60: 218–237.

29. Mochlyrosen, D., Khaner, H., and Lopez, J. (1991) Identification of intracellular receptor proteins for activated protein kinase C. Proceedings of the National Academy of Sciences of the United States of America 88: 3997–4000.

30. Neves, S.R., Ram, P.T., and Iyengar, R. (2002) G protein pathways. Science 296: 1636–1639.

31. Nishimura, M., Park, G., and Xu, J.R. (2003) The G-beta subunit *MGB1* is involved in regulating multiple steps of infection-related morphogenesis in *Magnaporthe grisea*. Molecular Microbiology 50: 231–243.

32. Nunez, A., Franco, A., Madrid, M., Soto, T., Vicente, J., Gacto, M., and Cansado, J. (2009) Role for RACK1 Orthologue Cpc2 in the Modulation of Stress Response in Fission Yeast. Molecular Biology of the Cell 20: 3996–4009.

33. Ortiz, C.S., and Shim, W.B. (2013) The role of MADS-box transcription factors in secondary metabolism and sexual development in the maize pathogen *Fusarium verticillioides*. Microbiology 159: 2259–2268.

34. Palmer, D.A., Thompson, J.K., Li, L., Prat, A., and Wang, P. (2006) Gib2, a novel Gβ-like/RACK1 homolog, functions as a Gβ subunit in cAMP signaling and is essential in *Cryptococcus neoformans*. Journal of Biological Chemistry 281: 32596–32605.

35. Rachfall, N., Schmitt, K., Bandau, S., Smolinski, N., Ehrenreich, A., Valerius, O., and Braus, G.H. (2013) RACK1/Asc1p, a Ribosomal Node in Cellular Signaling. Molecular & Cellular Proteomics 12: 87–105.

36. Ramanujam, R., Calvert, M.E., Selvaraj, P., and Naqvi, N.I. (2013) The late endosomal HOPS complex anchors active G-protein signaling essential for pathogenesis in *Magnaporthe oryzae*. PLoS Pathog 9: e1003527.

37. Ron, D., Chen, C.H., Caldwell, J., Jamieson, L., Orr, E., and Mochly-Rosen, D. (1994) Cloning of an intracellular receptor for protein kinase C: a homolog of the *β* subunit of G proteins. Proc Natl Acad Sci U S A 91: 839–843.

38. Rosen, S., Yu, J.H., and Adams, T.H. (1999) The *Aspergillus nidulans sfaD* gene encodes a G protein *β* subunit that is required for normal growth and repression of sporulation. EMBO J 18: 5592–5600.

39. Sagaram, U.S., and Shim, W.B. (2007) *Fusarium verticillioides GBB1*, a gene encoding heterotrimeric G protein *β* subunit, is associated with fumonisin B biosynthesis and hyphal development but not with fungal virulence. Mol Plant Pathol 8: 375–384.

40. Schneider, C.A., Rasband, W.S., and Eliceiri, K.W. (2012) NIH Image to ImageJ: 25 years of image analysis. Nat Methods 9: 671–675.

41. Schultzhaus, Z., Yan, H., and Shaw, B.D. (2015) *Aspergillus nidulans* flippase DnfA is cargo of the endocytic collar and plays complementary roles in growth and phosphatidylserine asymmetry with another flippase, DnfB. Mol Microbiol 97: 18–32.

42. Shim, W.B., and Woloshuk, C.P. (1999) Nitrogen repression of fumonisin B1 biosynthesis in *Gibberella fujikuroi*. FEMS Microbiol Lett 177: 109–116.

43. Shim, W.B., Sagaram, U.S., Choi, Y.E., So, J., Wilkinson, H.H., and Lee, Y.W. (2006) *FSR1* is essential for virulence and female fertility in *Fusarium verticillioides* and *F. graminearum*. Molecular Plant-Microbe Interactions 19: 725–733.

44. Smith, T.F., Gaitatzes, C., Saxena, K., and Neer, E.J. (1999) The WD repeat: a common architecture for diverse functions. Trends Biochem Sci 24: 181–185.

45. Tang, G., Chen, Y., Xu, J.R., Kistler, H.C., and Ma, Z. (2018) The fungal myosin I is essential for *Fusarium* toxisome formation. PLoS Pathog 14: e1006827.

46. Wang, C., Dong, X., Han, L., Su, X.D., Zhang, Z., Li, J., and Song, J. (2016) Identification of WD40 repeats by secondary structure-aided profile-profile alignment. J Theor Biol 398: 122–129.

47. Wang, P., Perfect, J.R., and Heitman, J. (2000) The G-protein *β* subunit *GPB1* is required for mating and haploid fruiting in *Cryptococcus neoformans*. Molecular and Cellular Biology 20: 352–362.

48. Wek, R.C., Staschke, K.A., and Narasimhan, J. (2004) 7 Regulation of the yeast general amino acid control pathway in response to nutrient stress. In *Nutrient-induced responses in eukaryotic cells*: Springer, pp. 171–199.

49. Xue, C., Hsueh, Y.-P., and Heitman, J. (2008) Magnificent seven: roles of G protein-coupled receptors in extracellular sensing in fungi. FEMS microbiology reviews 32: 1010–1032.

50. Yan, H., Huang, J., Zhang, H., and Shim, W.B. (2019) A Rab GTPase protein FvSec4 is necessary for fumonisin B1 biosynthesis and virulence in *Fusarium verticillioides*. Curr Genet.

51. Yin, Z., Zhang, X., Wang, J., Yang, L., Feng, W., Chen, C. et al. (2018) MoMip11, a MoRgs7-interacting protein, functions as a scaffolding protein to regulate cAMP signaling and pathogenicity in the rice blast fungus *Magnaporthe oryzae*. Environ Microbiol.

52. Yu, H.Y., Seo, J.A., Kim, J.E., Han, K.H., Shim, W.B., Yun, S.H., and Lee, Y.W. (2008) Functional analyses of heterotrimeric G protein Gα and Gβ subunits in *Gibberella zeae*. Microbiology-Sgm 154: 392–401.

53. Zeller, C.E., Parnell, S.C., and Dohlman, H.G. (2007) The RACK1 ortholog Asc1 functions as a G-protein *β* subunit coupled to glucose responsiveness in yeast. Journal of Biological Chemistry 282: 25168–25176.

54. Zhang, D., Wang, Y., Shen, J., Yin, J., Li, D., Gao, Y. et al. (2018a) *OsRACK1A*, encodes a circadian clock-regulated WD40 protein, negatively affect salt tolerance in rice. Rice (N Y) 11: 45.

55. Zhang, H., Mukherjee, M., Kim, J.E., Yu, W., and Shim, W.B. (2018b) Fsr1, a striatin homologue, forms an endomembrane-associated complex that regulates virulence in the maize pathogen *Fusarium verticillioides*. Mol Plant Pathol 19: 812–826.

56. Zhang, X., Jain, R., and Li, G. (2016) Roles of Rack1 Proteins in Fungal Pathogenesis. Biomed Res Int 2016: 4130376.

