## Supplementary Figures for "Characterization of non-canonical G beta-like protein FvGbb2 and its relationship with heterotrimeric G proteins in *Fusarium verticillioides*"

#### Supplementary Figure Legends

**Fig. S1. Sequence alignment and domain analysis of Gβ-like proteins in fungal species.** We aligned amino acid sequences of *Saccharomyces. cerevisiae* ScAsc1, *Fusarium verticillioides* FvGbb2, *F. verticillioides* FvGbb1, *Magnaporthe oryzae* MoMip11, *Neurospora crassa* NcGnb1, and *Cryptococcus neoformans* CnGib2. Identical and similar sequences were indicated in white characters with black backgrounds and black characters in a white box, respectively. FvGbb2 is predicted to have 28 beta strands. ScAsc1 protein structure was from PDB (ID:3FRX) (Coyle et al., 2009).

**Fig. S2. Schematic description of split marker approach for generating gene deletion mutants ΔFvgbb2, ΔFvgbb1, and ΔFvgpa2 in *F. verticillioides.*** (A) Hygromycin gene (*HYG*) was used to replace the *FvGBB2* gene and generate ΔFvgbb2 mutant. (B) Geneticin gene (*GEN*) was used to replace the *FvGBB1 gene* and generate ΔFvgbb1 mutant. (C) Geneticin gene (*GEN*) was used to replace the *FvGPA2 gene* and generate ΔFvgpa2 mutant. (D) Total RNA samples were extracted from mycelia from 7-day myro liquid culture. Mutants ΔFvgbb2, ΔFvgbb2-gbb1, ΔFvgbb2-gpa2 were subject to qPCR using *FvGBB2* gene primer. *FvGBB2* transcript was not detectable in ΔFvgbb2, ΔFvgbb2-gbb1, ΔFvgbb2-gpa2 mutants when compared to the wild-type (WT) progenitor. (E) *FvGBB1* gene expression was assayed in WT, ΔFvgbb1, and ΔFvgbb2-gbb1. (F) *FvGPA2* gene expression was assayed in WT, ΔFvgpa2, and ΔFvgbb2-gpa2.

**Fig. S3. Perithecia formation inf wild-type (WT) and deletion mutant strains.** (A) Expression levels of conidia related genes in ΔFvgbb2 were compared to the WT in YEPD liquid medium. (B)

(B) WT, ΔFvgbb2, ΔFvgbb1, ΔFvgpa2, ΔFvgbb2-gbb1and ΔFvgbb2-gpa2 strains, which all have MAT-1 mating type, were crossed to *F. verticillioides* m3120 strain (MAT-2) in carrot medium. After 3-week incubation, perithecia formation was observed.

**Fig. S4. *Fusarium verticillioides* colonization in autoclaved cracked kernels and surface sterilized kernels.** Wild-type (WT), ΔFvgbb2, ΔFvgbb1, ΔFvgpa2, ΔFvgbb2-gbb1, ΔFvgbb2-gpa2 and complementation strains were cultured in (A) 2-g cracked autoclaved (non-viable) kernels and (B) four sterilized surface-sterilized (viable) kernels after 8 days of incubation at room temperature. (C) Pigment production when strains are grown in myro liquid medium. Two mutants ΔFvgbb2 and ΔFvgbb2-gbb1 exhibited dramatic alteration in pigmentation when cultured for 7 days with shaking.

**Fig. S5. FvGbb2 protein localization assays.** (A) Schematic representation of FvGbb2-GFP, mCherry-FvGbb1, FvGpa2-mCherry constructs. (B) FvGbb2-GFP was expressed in the ΔFvgbb2 mutant in (B) FB1 non-inducing 0.2xPDB medium and in (C) FB1-inducing myro medium. Bar = 5 μm. (D) FvGbb2-GFP was expressed in the WT and with 10 μM FM4-64 stain after a 30-min incubation. Bar = 5 μm.

**Fig. S6. Interaction between FvGbb2 and MAPK kinase cascade proteins. (A)** FvGbb2 did not show interaction with three MAPK cascade proteins when tested by yeast two-hybrid assay. FvGbb2 was co-transformed with FvBck1, FvMkk1/2 or FvSlt2 into AH109 strain. pGBKT7–53 and pGADT7-T were used as a positive control. pGBKT7–Lam and pGADT7-T were used as a negative control. (B) The split luciferase complementation analysis was performed to confirm yeast two-hybrid assays. FvGbb1-FvGpa2 complementation was used as a positive control.

### Yan & Shim. Fig. S1

```
ScAsc1                                                            1
ScAsc1   ......................................................MASNEV
FvGbb2   .........................................................MAEQ
FvGbb1   MNSQGNNDVSPEAMQSRIQQARREAETLKDRIKRKKDDLADTTLRAVAQQAHEPIPKNQL
NcCpc2   .........................................................MAEQ
MoMip11  .........................................................MAEQ
CnGib2   .........................................................MAEH
```

```
         β1          β2             β3          β4            β5
ScAsc1                        TT        TT               TT
           10        20        30        40        50        60
ScAsc1   LVLRGTLEGHNGWVTSLATSAGQPNLLLSASRDKTLISWKLTGDDQKFGVPVRSFKGHSH
FvGbb2   LILKGTLEGHNGWVTSLATSMENPNMLLSASRDKTLIIWNLTRDETQYGYPKRSLHGHSH
FvGbb1   MKAKRTLKGHLAKIYAMHWSTDRRHLV.SASQDGKLIIWDAYTTNKVHAIPLR.....SS
NcCpc2   LILKGTLEGHNGWVTSLATSMENPNMLLSSSRDKSLIIWNLTRDETSYGYPKRRLHGHSH
MoMip11  LILKGTLEGHNGWVTSLATSMENPNMLLSSSRDKTLIIWNLTRDETSYGYPKRSLKGHSH
CnGib2   LMFKGNLAGHNGWVTAIATSSENPDMILTASRDKTVIAWQLTREDNLYGFPKKILHGHNH
```

```
          β6          β7        β8                  β9         β10
ScAsc1             TT       TT                                    TT
           70        80        90        100       110       120
ScAsc1   IVQDCTLTADGAYALSASWDKTLRLWDVAT...GET..YQRFVGHKSDVMSVDIDKKASM
FvGbb2   IVSDCVISSDGAYALSASWDKTLRLWELAS...GTT..TRRFVGHTNDVLSVSFSADNRQ
FvGbb1   WVMTCAYAPSGNFVACGGLDNICSIYNLNQQRDGPTRVARELSGHAGYLSCCRF.INDRS
NcCpc2   IVSDCVISSDGAYALSASWDKTLRLWELST...GTT..TRRFVGHTNDVLSVSFSADNRQ
MoMip11  IVSDCVISSDGAYALSASWDKTLRLWELAT...GTT..TRRFVGHTNDVLSVSFSADNRQ
CnGib2   FVSDVAISSDGQFALSSSWDHTLRLWDLNT...GLT..TKKFVGHTGDVLSVSFSADNRQ
```

```
         β11         β12         β13           β14              β15
ScAsc1        TT          TT                                      TT
                   130       140       150       160       170
ScAsc1   IISGSRDKTIKVWTIKGQCLATLL..GHNDWVSQVRVVPNEQADDDSVTIISAGNDKMVK
FvGbb2   IVSGSRDRTIKLWNTLGDCKYTITEKGHTEWASCVRFSPNPQNP....VIVSAGWDKLVK
FvGbb1   ILTSSGDMTCMKWDIETGQKVTEFADHLGDVMSISLNPTNQN......TFISGACDAFAK
NcCpc2   IVSGSRDRTIKLWNTLGDCKFTITEKGHTEWVSCVRFSPNPQNP....VIVSSGWDKLVK
MoMip11  IVSGSRDRSIKLWNTLGDCKYTITEKGHSEWVSCVRFSPNPQNP....VIVSSGWDKLVK
CnGib2   IVSASRDRSIKLWNTLGECKFDIVEDGHTEWVSCVRFSPNPALP....VIISAGWDKTVK
```

```
         β16       β17           β18           β19       β20       β21
ScAsc1                                    TT          TT
         180       190       200       210       220       230
ScAsc1   AWNLNQFQIEADFIGHNSNINTLTASPDGTLIASAGKDGEIMLWNLAAKKAMYTLSAQDE
FvGbb2   VWELSTCKLQTDHIGHTGYINTVTISPDGSLCASGGKDGTTMLWDLNESKHLYSLNANDE
FvGbb1   LWDIRAGKAVQTFAGHESDINAIQFFPDGHSFVTGSDDATCRLFDIRADRELNLYGSESI
NcCpc2   VWELSSCKLQTDHIGHTGYINAVTISPDGSLCASGGKDGTTMLWDLNESKHLYSLNANDE
MoMip11  VWELSSCKLQTDHIGHTGYINTVTISPDGSLCASGGKDGTTMLWDLNESKHLYSLNANDE
CnGib2   VWELSNCKLKTTHHGHTGYLNTLAVSPDGSLAASGGKDGITMLWDLNEGKHLYSLDAGDV
```

```
          β22                β23      β24        β25          η1     β26
ScAsc1                          TT                      TT
         240       250       260       270       280       290
ScAsc1   VFSLAFSPN....RYWLAAATATGIKVFSLDPQYLVDDLRPEFAGYS.KAAEPHAVSLAW
FvGbb2   IHALVFSPN....RYWLCAATASSIIIFDLEKKSKVDELKPEFPAVGKKSREPECVSLAW
FvGbb1   LCGITSVATSVSGRLLFAGYDDFECKVWDITRGEKVGSL.........VGHENRVSCLGV
NcCpc2   IHALVFSPN....RYWLCAATSSSIIIFDLEKKSKVDELKPEFQNIGKKSREPECVSLAW
MoMip11  IHALVFSPN....RYWLCAATASSIIIFDLEKKSKVDELKPEFAAVGKKSREPECISLAW
CnGib2   INALVFSPN....RYWLCAATASSIKIFDLESKSLVDDLQPDFDGLSDKARKPECTSLAW
```

```
                  β27          β28
ScAsc1    TT            TT
             300       310
ScAsc1   SADGQTLFAGYTDNVIRVWQVMTAN
FvGbb2   SADGQTLFAGYTDNIIRAWGVMSRA
FvGbb1   SNDGTSLCTGSWDSLLKIWAY....
NcCpc2   SADGQTLFAGYTDNIIRAWGVMSRA
MoMip11  SADGQTLFAGYTDNIIRAWGVMSRA
CnGib2   SADGQTLFAGFSDNLVRVWAVVA..
```

Yan & Shim. Fig. S2

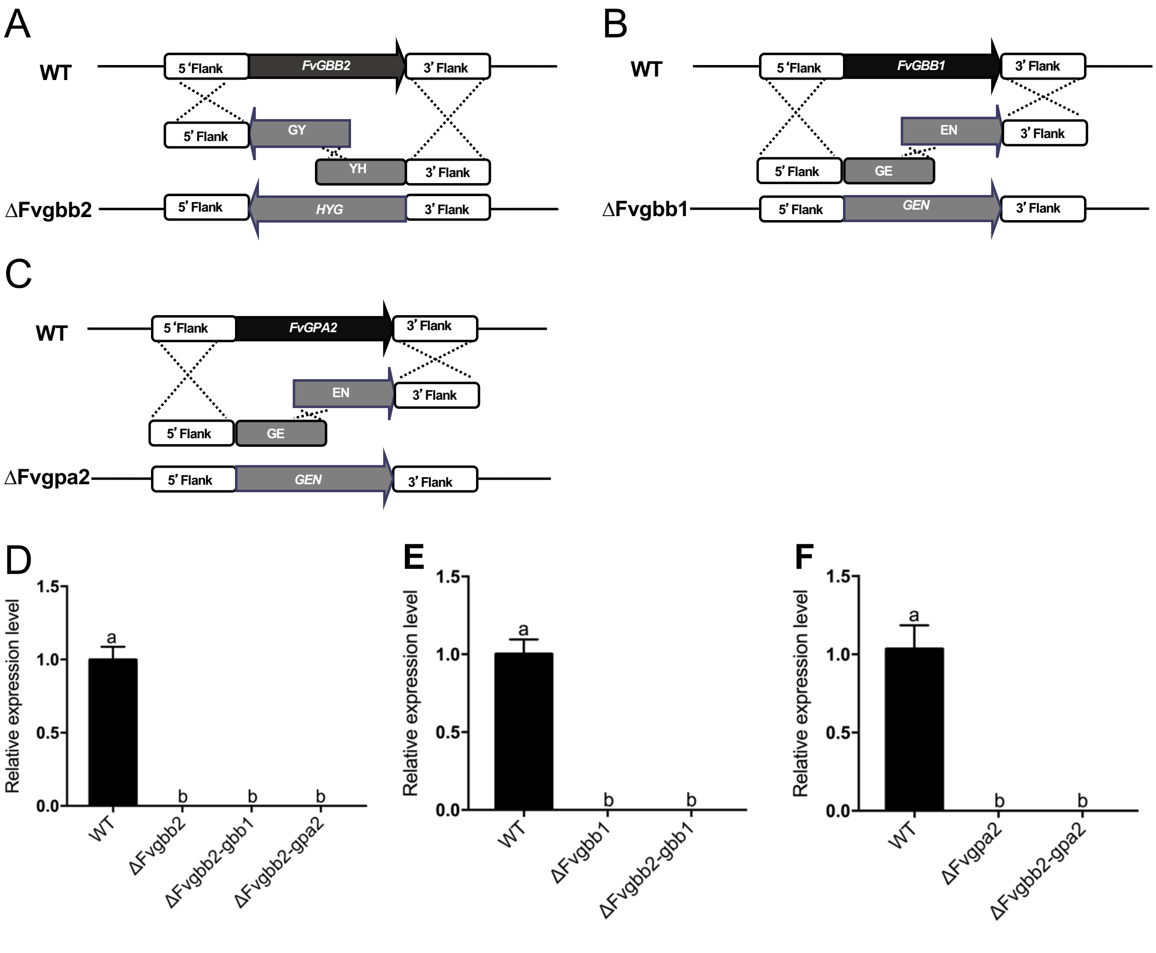

Yan & Shim. Fig. S3

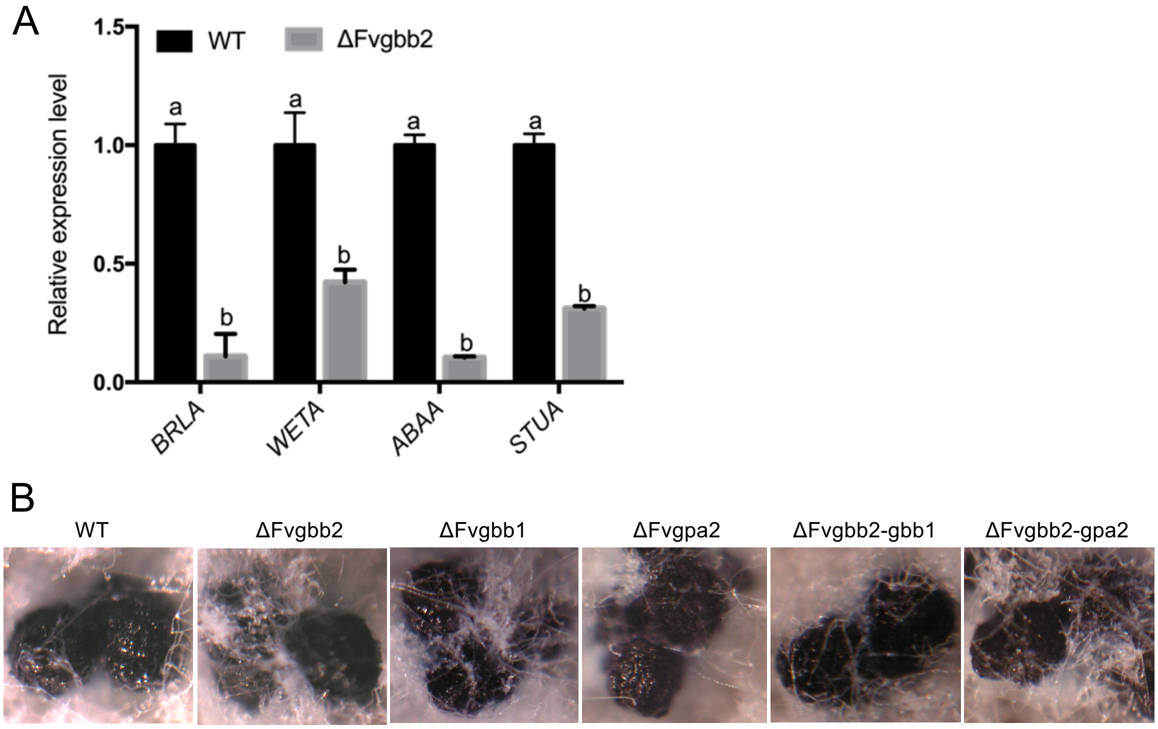

Yan & Shim. Fig. S4

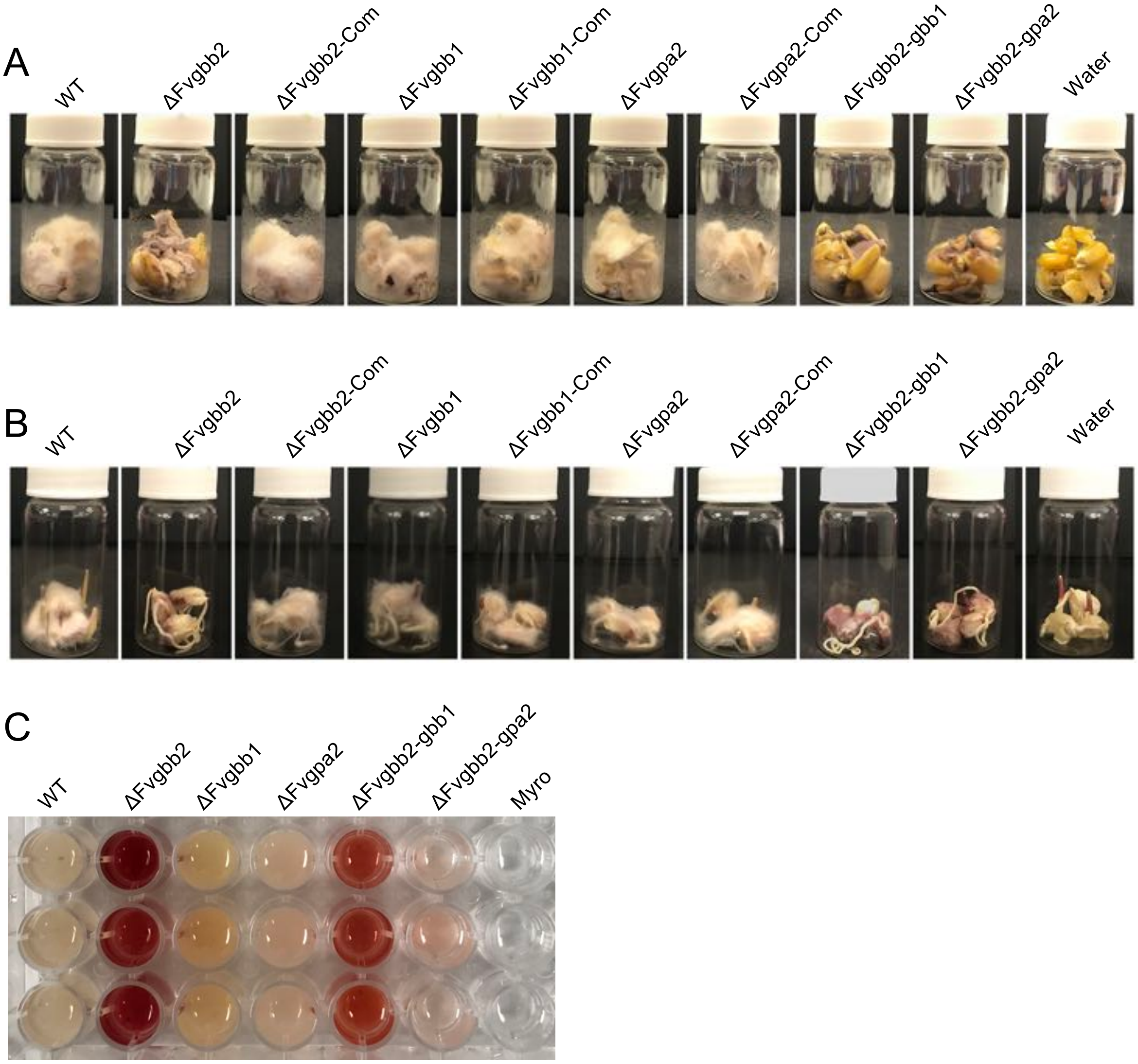

Yan & Shim. Fig. S5

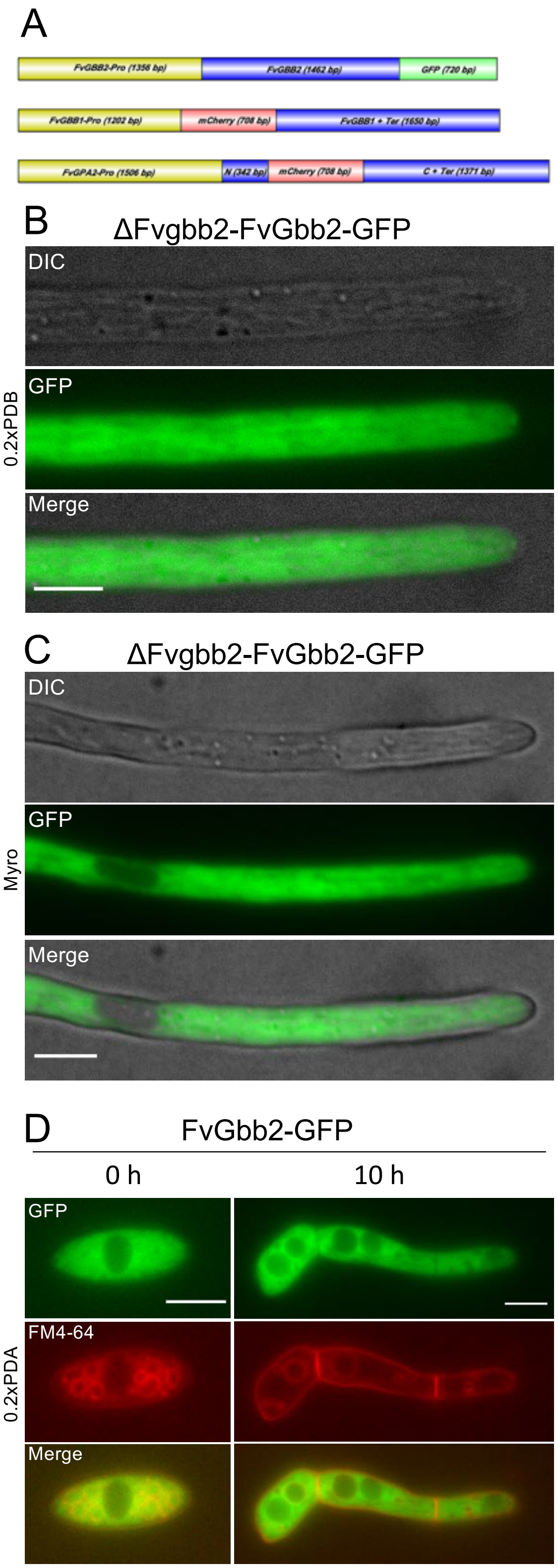

Yan & Shim. Fig. S6

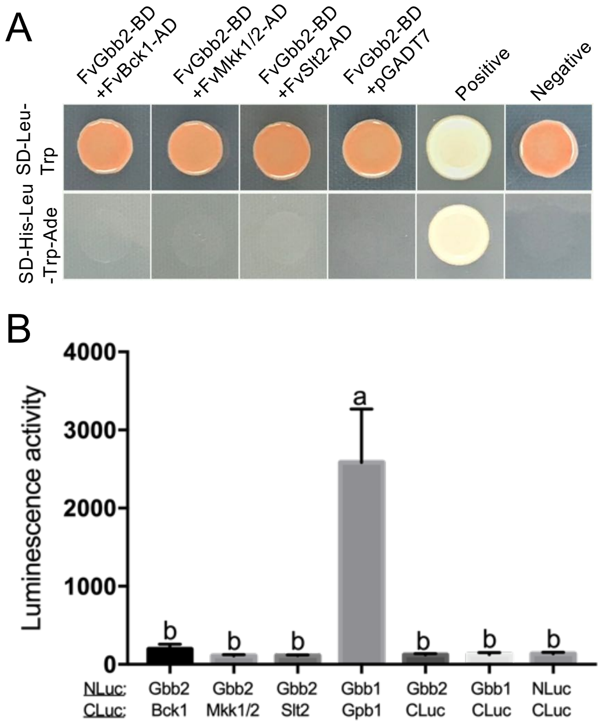
