## Supplementary Tables for "Characterization of non-canonical G beta-like protein FvGbb2 and its relationship with heterotrimeric G proteins in *Fusarium verticillioides*"

### Table S1. Relative mRNA expression level of 15 PKS gene in deletion mutant strains

| | **Relative transcriptional level** | | | | | |
|---|---|---|---|---|---|---|
| **Gene** | WT | ΔFvgbb2 | ΔFvgbb1 | ΔFvgpa2 | ΔFvgbb2-gbb1 | ΔFvgbb2-gpa2 |
| *PKS1* | 1[a] | 0.24 ± 0.01 | 0.45 ± 0.19 | 0.58 ± 0.14 | 0.10 ± 0.09 | 0.28 ± 0.13 |
| *PKS2* | 1 | ND | 0.31 ± 0.15 | 0.62 ± 0.28 | 1.16 ± 0.46 | 0.46 ± 0.13 |
| *PKS3* | 1 | 762.42 ±168 | 1.89 ± 0.06 | 6.78 ± 2 79 | 0.47 ± 0.16 | 23.37 ± 10.26 |
| *PKS4* | 1 | ND | 1.28 ± 0.12 | 1.64 ± 0.67 | ND | ND |
| *PKS5* | 1 | 0.25 ± 0.01 | 0.86 ± 0.19 | 0.99 ± 0.01 | 0.75 ± 0.15 | ND |
| *PKS6* | 1 | 208.70 ± 28.52 | 1.32 ± 1.17 | 12.05 ± 4.15 | 131.84 ± 42.62 | 54.22 ± 22.75 |
| *PKS7* | 1 | 0.48 ± 0.21 | 0.65 ± 0.26 | 0.83 ± 0.42 | 1.14 ± 0.50 | 0.59 ± 0.15 |
| *PKS8* | 1 | 0.10 ± 0.02 | 0.23 ± 0.10 | 0.31 ± 0.20 | 0.06 ± 0.04 | 0.72 ± 0.18 |
| *PKS9* | 1 | 0.003 ± 0.001 | 0.82 ± 0.10 | 1.50 ± 0.10 | ND | 0.08 ± 0.01 |
| *PKS10* | 1 | 0.02 ±0.001 | 3.53 ± 1.19 | 1.51 ± 0.44 | 0.29 ± 0.01 | 0.08 ± 0.02 |
| *PKS11* | 1 | ND | ND | 0.86 ± 0.34 | ND | ND |
| *PKS12* | ND | ND | ND | ND | ND | ND |
| *PKS13* | 1 | 0.42 ± 0.12 | 0.95 ± 0.12 | 1.44 ± 0.55 | 2.46 ± 0.69 | 0.58 ± 0.14 |
| *PKS14* | 1 | 1.36 ± 0.15 | 0.76 ± 0.20 | ND | 0.55 ± 0.24 | 0.55 ± 0.02 |
| *PKS15* | 1 | 0.15 ± 0.02 | 0.17 ± 0.04 | 0.23 ± 0.10 | 0.43 ± 0.14 | 5.48 ± 0.56 |

Strains mycelium were harvested from 7 days of incubation on myro liquid medium. Then, total RNA samples were extracted. qPCR analysis of gene expressional levels on was conducted with SYBR-Green. Each gene expression was normalized to β-tubulin gene (FVEG_04081) expression level. Gene expressions were calculated using $2^{-\Delta\Delta Ct}$. Wild-type expression level was standardized to 1.0. Three replicates were performed for each test. *PKS5*, *PKS14*, *PKS15* expression levels were very low.

**Table S2.** Primers used in this study

| Primer | Primer sequence (5'-3') | Application |
|---|---|---|
| Gbb2_LF.F2 | GCT CAG TGG ACA ACC AGA CAA A | validation of *FvGBB2* 5' deletion |
| Gbb2_LF.F1 | TCC TTC TTC CTT TCC CGA CTT TC | amplify *FvGBB2* 5' flank sequence |
| Gbb2_LF.F/N | CGC AAG CAG AAA GAG GTC AAG AA | nested primer for amplify *FvGBB2-LF + YG* |
| Gbb2_LF.R1 | TAG ATG CCG ACC GGG AAC GTT TGC GGT GAC GAG GCG AT | amplify *FvGBB2* 5' flank sequence |
| Gbb2_RF.F1 | CCA CTA GCT CCA GCC AAG ATT GCT CTA CCC TAA AAA AGA GGC | amplify *FvGBB2* 3' flank sequence |
| Gbb2_RF.RN | TCT CTT ATT CAC CAG CGG ACA | nested primer for amplify *FvGBB2-RF + HY* |
| Gbb2_RF.R1 | GTT GGG CGA TCA CCA TAA CTA | amplify *FvGBB2* 3' flank sequence |
| Gbb1/LF/F2 | GTG TTT GGG TGA GCG GAT G | validation of *FvGBB1* 5' deletion |
| Gbb1-LF/F1 | CGG GTA CTT GAT GTT TGG TGG A | amplify *FvGBB1* 5' flank sequence |
| Gbb1-LF/FN | ACG GAT GAA CTG GAG TCG A | nested primer for amplify *FvGBB1-LF + GE* |
| Gbb1-LF/R1 | GGC GTT ACC CAA CTT AAT CGA TCG ACC AGT CAA CTT TTG TTG T | amplify *FvGBB1* 5' flank sequence |
| Gbb1_RF/F1 | TTC CAC ACA ACA TAC GAG CCT TGT TCT TTG CGA CTA TGA ACC TT | amplify *FvGBB1* 3' flank sequence |
| Gbb1_RF/RN | ACA CAC GTC CGA CAG CAA ATA | nested primer for amplify *FvGBB1-RF + EN* |
| Gbb1_RF/R1 | ACG GGA TAA GTG CGG AAT GTT | amplify *FvGBB1* 3' flank sequence |
| Gpa2_ LF/F2 | ACA TGG AGG CTT GTG AGG TTA T | validation of *FvGPA2* 5' deletion |
| Gpa2_LF/F1 | ACC TTA CTT TCC CGT CCC TC | amplify *FvGPA2* 5' flank sequence |
| Gpa2_LF/FN | TTT GGC ATT ACT TCC GCT TGC | nested primer for amplify *FvGPA2-LF + GE* |
| Gpa2_LF/R1 | GGC GTT ACC CAA CTT AAT CGT GTG GCG GAT CAC AAA ATA GCT | amplify *FvGPA2*5' flank sequence |
| Gpa2_ RF/F1 | TTC CAC ACA ACA TAC GAG CCT AGC GAA TCA CTG CTC ATT TGA | amplify *FvGPA2*3' flank sequence |
| Gpa2_ RF/RN | GTC ATT TAC GCC AAG CGA CTA T | nested primer for amplify *FvGPA2-RF + EN* |
| Gpa2_RF/R1 | AAG CTC CAA TCC ACA TCC AGA | amplify *FvGPA2* 3' flank sequence |
| HYG/F | TTG GCT GGA GCT AGT GGA GGT CAA | amplify HY fragment |

| | | |
|---|---|---|
| HY/R | GTA TTG ACC GAT TCC TTG CGG TCC GAA | amplify HY fragment |
| HYG/R | GTT CCC GGT CGG CAT CTA CTC TAT | amplify YG fragment |
| YG/F | GAT GTA GGA GGG CGT GGA TAT GTC CT | amplify YG fragment |
| GEN/F | CGA TTA AGT TGG GTA ACG CCA G | amplify GE fragment |
| GE/R | ATC ACG GGT AGC CAA CGC TA | amplify GE fragment |
| EN/F | TCG ACC ACC AAG CGA AAC AT | amplify EN fragment |
| GEN/R | GGC TCG TAT GTT GTG TGG AAT T | amplify EN fragment |
| Gbb2-BD-F | CCG GAA TTC ATG GCC GAA CAA TTG ATC CTG AA | construction of pGBDT7-*FvGBB2* |
| Gbb2-BD-R | CCG GGA TCC TTA TGC CCT CGA CAT GAC ACC | construction of pGBDT7 -*FvGBB2* |
| Gpa1-AD-F | CCG GAA TTC ATG GGT TGC GGA ATG AGC | construction of pGADT7-*FvGPA1* |
| Gpa1-AD-R | CCG GGA TCC TTAGAT GAG ACC ACA GAG ACG CAG | construction of pGADT7 -*FvGPA1* |
| Gpa2-AD-F | CCG GAA TTC ATG GGC GCA TGC ATG AGC T | construction of pGADT7-*FvGPA2* |
| Gpa2-AD-R | CCG GGA TCC TCA AAG AAT GCC CGA GTC CTT AAG TG | construction of pGADT7 -*FvGPA2* |
| Gpa3-AD-F | CCG GAA TTC ATG CTG CAG AAA CAC ATG GCC | construction of pGADT7-FvGPA3 |
| Gpa3-AD-R | CCG GGA TCC TTA GAG AAT GAG CTG CTT GAG GTT G | construction of pGADT7 -*FvGPA3* |
| Gpb1-AD-F | CCG GAA TTC ATG CCT CAG TAC ACT TCT CGC | construction of pGADT7-*FvGPB1* |
| Gpb1-AD-R | CCG GGA TCC TTA CAT GAC CAC ACA GCA GC | construction of pGADT7 -*FvGPB1* |
| Mkk1/2-AD-F | CGG AAT TCA TGG CCG ACC AAC AGC CT | construction of pGADT7-*FvMKK1/2* |
| Mkk1/2 -AD-R | TCC CCC GGG CTA AGA GTC CTT AGG TTG TTC ATC CC | construction of pGADT7 -*FvMKK1/2* |
| Slt2_AD-F | CGG AAT TCA TGT CGG ACC TTC AAG GAC G | construction of pGADT7-*FvSLT2* |
| Slt2_AD-R | CGG GAT CCT TAT CTC CTG GAG GCA TCC A | construction of pGADT7 -*FvSLT2* |
| Bck1-AD-F1 | ACCAGATTACGCTCATATG ATGAAGGGTAGCGATAGCTTG | construction of pGADT7 -*FvBCK1* |
| Bck1-AD-R1 | ATT TGG CGA ATC CAT GGG ATC | construction of pGADT7-*FvBCK1* |
| Bck1-AD-F2 | TCCCATGGATTCGCCAAATACACCTCTCAATGCCCCTG | construction of pGADT7 -*FvBCK1* |
| Bck1-AD-R2 | CTA TTT CGT TGG ACA TCC TCG G | construction of pGADT7-*FvBCK1* |
| Bck1-AD-F3 | CCG AGG ATG TCC AAC GAA ATA G | construction of pGADT7 -*FvBCK1* |
| Bck1-AD-R3 | CAG CTC GAG CTC GAT GGATCC TTA TTT GAA CGA CTT GAT TTT GTG ATA CAG | construction of pGADT7 -*FvBCK1* |
| Gbb2_nLUC/F | AAG CTC GAG TAG TCG ACA TGG CCG ACC AAC AGC CTG AA | construction of pFNLuc *FvGBB2* |

| | | |
|---|---|---|
| Gbb2_nLUC/R | CGT ACG AGA TCT GGT CGA CTG CCC TCG ACA TGA CAC CCCA | construction of pFNLuc *FvGBB2* |
| Gbb1_nLUC/F | AAG CTC GAG TAG TCG ACA TGA ACT CTC AAG GCA ACA ACG A | construction of pFNLuc-*FvGBB1* |
| Gbb1_nLUC/R | CGT ACG AGA TCT GGT CGA CGT AGG CCC AGA TTT TAA GCA G | construction of pFNLuc-*FvGBB1* |
| Gpa1_cLUC/F | CGT CCC GGG GCG GTA CCG GTT GCG GAA TGA GCA CAG A | construction of pFCLuc-*FvGPA1* |
| Gpa1_cLUC/R | TTG GAT CCC CGG GTA CCT TAG ATG AGA CCA CAG AGA CGC AG | construction of pFCLuc-*FvGPA1* |
| Gpa2_cLUC/F | CGT CCC GGG GCG GTA CCG GCG CAT GCA TGA GCT CG | construction of pFCLuc-*FvGPA2* |
| Gpa2_cLUC/R | TTG GAT CCC CGG GTA CCT CAA AGA ATG CCC GAG TCC TTA AGT G | construction of pFCLuc-*FvGPA2* |
| Gpa3_cLUC/F | CGT CCC GGG GCG GTA CCC TGC AGA AAC ACA TGG CCA | construction of pFCLuc-*FvGPA3* |
| Gpa3_cLUC/R | TTG GAT CCC CGG GTA CCT TAG AGA ATG AGC TGC TTG AGG TTG | construction of pFCLuc-*FvGPA3* |
| Gbb1_cLUC/F | CGT CCC GGG GCG GTA CCA ACT CTC AAG GCA ACA ACG A | construction of pFCLuc-*FvGBB1* |
| Gbb1_cLUC/R | TTG GAT CCC CGG GTA CCT TAG TAG GCC CAG ATT TTA AGC AG | construction of pFCLuc-*FvGBB1* |
| MKK1/2_cLUC/F | CGT CCC GGG GCG GTA CCG CCG ACC AAC AGC CTC AAA GT | construction of pFCLuc-*FvMKK1/2* |
| MKK1/2_cLUC/R | TTG GAT CCC CGG GTA CCC TAA GAG TCC TTA GGT TGT TCA TCC CAG | construction of pFCLuc-*FvMKK1/2* |
| Slt2_cLUC/F | CGT CCC GGG GCG GTA CCT CGG ACC TTC AAG GAC GGA | construction of pFCLuc-*FvSLT2* |
| Slt2_cLUC/R | TTG GAT CCC CGG GTA CCT TAT CTC CTG GAG GCA TCC A | construction of pFCLuc-*FvSLT2* |
| Bck1_cLUC/F | CGTCCCGGGGCGGTACC AAGGGTAGCGATAGCTTGC | construction of pFCLuc-*FvBCK1* |
| Bck1_cLUC/R | TTGGATCCCCGGGTACC TTA TTT GAA CGA CTT GAT TTT GTG ATA CAG | construction of pFCLuc-*FvBCK1* |
| cLUC_F: | GAT TGA CAA GGA TGG ATG GCT AC | screening paired with each gene cLUC/R |
| nLUC-R | GGT AGA TGA GAT GTG ACG AAC GTG | screening paired with each gene nLUC/F |

| | | |
|---|---|---|
| Gbb2-GFP-F | AGG GAA CAA AAG CTG GGT ACC CGC TAC GAC GGC ACA ATG TT | construction of pKNTG-*FvGBB2* |
| Gbb2-GFP-R | GCC CTT GCT CAC CAT AAG CTT TGC CCT CGA CAT GAC ACC | construction of pKNTG-*FvGBB2* |
| Gbb1-pro-F | AGG GAA CAA AAG CTG GGT ACC GTG TTT GGG TGA GCG GAT G | amplify *FvGBB1* native promoter |
| Gbb1-pro-R | CTC CTC GCC CTT GCT CAC CAT CGT GAT CGA CCA GTC AAC TTTT | amplify *FvGBB1* native promoter |
| Gbb1-F | GGC ATG GAC GAG CTG TAC AAG ATG AAC TCT CAA GGC AAC AAC GA | amplify *FvGBB1* coding sequence |
| Gbb1-R | TCA GTA ACG TTA AGT GGA TCC GTA TAG CCG TCA ACT CCT ATG GA | amplify *FvGBB1* coding sequence |
| Gpa2-pro-F | AGG GAA CAA AAG CTG GGT ACC ACA TGG AGG CTT GTG AGG TTA T | amplify *FvGPA2* native promoter |
| Gpa2-pro-R | TGA TCC GCC TCC TCA TTG CT | amplify *FvGPA2* native promoter |
| Gpa2-N-F | AGCAATGAGGAGGCGGATCA | amplify *FvGPA2* partial coding sequence |
| Gpa2-N- R | CTC CTC GCC CTT GCT CAC CAT GGC TTG GTA TTC TAA TAG GAA CTC C | amplify *FvGPA2* partial coding sequence |
| Gpa2-C-F | GGC ATG GAC GAG CTG TAC AAG GAG TCT GGC CCT CAA GCG CAA AT | amplify *FvGPA2* partial coding sequence and terminal |
| Gpa2-Ter-R | TCA GTA ACG TTA AGT GGA TCC AAG GGA AGC CAT ACG CCT GTT CTC | amplify *FvGPA2* partial coding sequence and terminal |
| mCherry-F | ATGGTGAGCAAGGGCGAGGAG | amplify mCherry fragment |
| mCherry-R | CTT GTA CAG CTC GTC CAT GCC | amplify mCherry fragment |
| Gbb2-qpcr_F1 | ATC GTC TCC GAC TGT GTC ATC TCC TCT GA | qPCR analysis |
| Gbb2-qpcr_R1 | TAG TAC CGC TGG CAA GCT CCC AGA | qPCR analysis |
| Gbb1-qPCR_F1 | TTC GTG GCT TGC GGT GGT CT | qPCR analysis |
| Gbb1-qPCR_R1 | TCA CGG GCA ACA CGG GTA GG | qPCR analysis |
| Gpa2-qPCR-F1: | TGA GCA GGT CTC CGC TAG GCA A | qPCR analysis |
| Gpa2-qPCR-R1 | TGT CGC TTG CGT CAA GTG GGG A | qPCR analysis |

| | | |
|---|---|---|
| Tub2-F | CAG CGT TCC TGA GTT GAC CCA ACA G | qPCR analysis |
| Tub2-R | CTG GAC GTT GCG CAT CTG ATC CTC G | qPCR analysis |
